# Structure-based analysis unveils co-origin of LPOR and nitrogenase-like proteins

**DOI:** 10.1101/2025.02.17.634645

**Authors:** Xizhe Sun, Lisong Ma, Fanli Zeng, Yiguo Hong, Ray Dixon, Zihe Rao, Guogang Zhao, Jin Zhao, Chao Zhang, Meng Wu, Chukang Ma, Xiaolan Yu, Ji Yang, Saul Purton, Elena Ermilova, Nigel Scrutton, Pedro Silva, Jianjun Zhao, Qi Cheng

## Abstract

Structures of nitrogenases, dark-operative protochlorophyllide oxidoreductases, and light-dependent protochlorophyllide oxidoreductases (LPOR) have been resolved. However, their evolutionary relatedness remains elusive. Here, we show, through structural alignment, that all subunits of nitrogenase-like proteins originated from a co-ancestral archaic one-subdomain precursor. LPOR evolved from the BchX/BchY subunits of nitrogenase-like chlorophyllide *a* oxidoreductase (COR), and the intermediary retinol dehydrogenase through possible genetic recombination. We thus establish previously unknown structural links among key enzymes involved in biological nitrogen-fixation (BNF) and photosynthesis, unraveling structure-guided functional evolution from a single-subunit iron protein to multi-subunit nitrogenase-like COR, and to the single-subunit LPOR for phototrophic metabolism *via* bacteriochlorophyll, retinal, and chlorophyll. This work also demonstrates structural similarities are imperative for inferring distant origins of functionally divergent proteins, particularly those lacking primary amino-acid sequence identity. Moreover, our findings coupled with AI may be exploited to design innovative light-driven CORs and/or light-utilizing nitrogenases with enhanced efficacy of photosynthesis and BNF.

**Graphical abstract:** 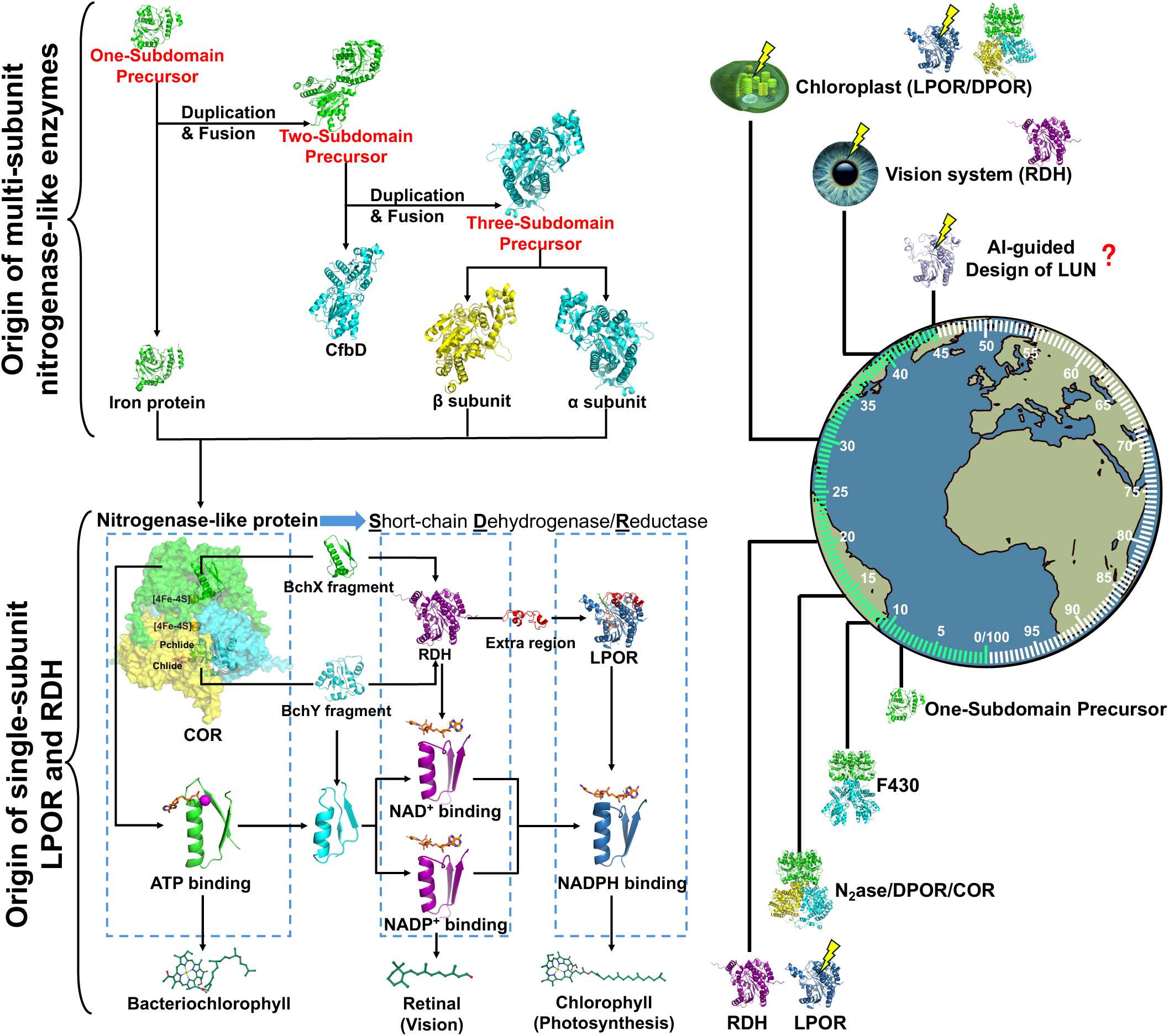

**Video abstract:** https://www.youtube.com/watch?v=b0qBE1JMaH0

**Highlights:**

- LPOR evolves from BchX and BchY subunits of the nitrogenase-like protein (NGLP) COR
- All subunits of NGLPs can be traced back to the ancestral one-subdomain precursor
- Retinol dehydrogenase acts as a crucial intermediate in the COR to LPOR evolution
- Structure-based design of LUN is showcased to revolutionize protein engineering

## INTRODUCTION

Photosynthesis and nitrogen-fixation are two of the most vital processes for sustaining life on Earth. Photosynthesis, which uses light energy to convert carbon dioxide and water into glucose and oxygen, played a vital role in the accumulation of oxygen during the Great Oxygenation Event (GOE) approximately 2.4 billion years ago^1,2^. This transformative process, driven by oxygenic cyanobacteria, was facilitated by the evolution of light-dependent protochlorophyllide oxidoreductase (LPOR), a key enzyme in chlorophyll biosynthesis that enabled efficient light harvesting and photosynthesis, ultimately contributing to the atmospheric oxygen buildup that reshaped Earth’s biosphere^2–5^. The light-dependent reactions of photosynthesis occur in the chloroplast thylakoid membrane and involve light absorbance and conversion into chemical energy by light-absorbing pigment chlorophylls^6,7^. Chlorophyll biosynthesis depends on LPOR and/or dark-operative protochlorophyllide oxidoreductase (DPOR), two key catalysts which catalyze the conversion (reduction) of Pchlide to chlorophyllide^8^. LPOR belongs to the short-chain dehydrogenase/reductase (SDR) family and consists of a single polypeptide^9^. Crystal structural analyses of cyanobacterial LPOR from *Thermosynechococcus elongatus* and *Synechocystis* sp. and modeling of the ternary Pchlide-NADPH-LPOR complex reveal that Pchlide binding, photosensitization, and photochemical conversion to chlorophyllide require multiple interactions between the LPOR active site and Pchlide^10^. Notably, by determining its atomic structure, AtPORB, one of the *Arabidopsis thaliana* LPOR isoforms, was found to coordinate chlorophyll biosynthesis and photosynthetic membrane biogenesis^8,11^. This was evidenced by the formation of LPOR oligomers that position Pchlide into helical filaments in the outer leaflet of the membrane, indicating that bifunctional LPOR is able to regulate chlorophyll biosynthesis, interacts with ferrochelatase, and acts as a “master switch” for photomorphogenesis in early plant development^8,11,12^.

Biological nitrogen (N_2_) fixation (BNF) emerged about 3 billion years ago prior to oxygenic photosynthesis. BNF involves nitrogenases that are two-component metalloproteins i.e., the iron (Fe) and molybdenum-iron (MoFe) proteins. These proteins are responsible for BNF by catalyzing the reduction of N_2_ to ammonia (NH_3_) using ATP and reducing power^13–16^. Among the three known homologous nitrogenase isoforms, molybdenum (Mo) nitrogenase, encoded by *nifHDK*, is the most widespread^17^. NifH from *Azotobacter vinelandii* forms a homodimer and undergoes conformational changes upon binding to MgATP^18–21^. During BNF, electron transfers to the MoFe protein (NifDK) with the Fe protein acting as the electron donor. This process is followed by ATP hydrolysis, which catalyzes the release of NifH from NifDK^22^. The heterotetramer MoFe protein contains “two α (NifD) and two β (NifK)” subunits, and the FeMo cofactor (FeMoco) is located in the α subunit. The metal P-cluster embraces eight iron and seven sulfur atoms and bridges the two subunits, thus acting as an intermediary in the electron transfer process from Fe protein to FeMoco^23,24^. The Fe-MoFe nitrogenase complex was initially proposed to comprise one Fe-protein dimer binding to each pair of αβ subunits in the MoFe-protein heterotetramer^25,26^. However, recent research indicates that the MoFe-protein binds only one Fe-protein at a time during turnover, indicative of half-reactivity^27^.

Nitrogenase-like enzymes include DPOR, chlorophyllide *a* oxidoreductase (COR) that catalyzes bacteriochlorophyll (BChl) biosynthesis, and an enzyme designated as F430 in this study involved in the synthesis of the porphyrin cofactor F430. This F430 enzyme is also known as the archaeal Ni^2+^-sirohydrochlorin a,c-diamide reductase, which is essential for the activity of methyl coenzyme-M reductase involved in methanogenesis and anaerobic methane oxidation^28–32^. These enzymes share a similar subunit arrangement within the protein complex and require ATP for substrate reduction as those in nitrogenase^28,31,33^. Nitrogenase-like protein catalysts usually comprise catalytic component 1 (like MoFe protein) and component 2 (like Fe protein), and are biologically active only upon the two components dock together^34^. For instance, components 1 and 2 of the DPOR complex contain a BchN/ChlN-BchB/ChlB heterotetramer and a BchL/ChlL homodimer, and act in concert to catalyze the reduction of Pchlide to Chlide coupled with ATP hydrolysis^35^. The BchL/ChlL subunit of DPOR shares about 50% amino acid sequence similarity with the nitrogenase Fe protein NifH^36^. Similarly, the BchN/ChlN and BchB/ChlB proteins share sequence and structure similarity with the NifD and NifK subunits of nitrogenase MoFe protein, respectively^33,36,37^. Interestingly, the Pchlide-binding pocket in the DPOR complex is in the same position as the FeMoco ligand binding site in the nitrogenase complex^33^. Despite DPOR resembles nitrogenase in primary amino-acid sequence and structure, the Fe protein NifH cannot be substituted by BchL to reduce nitrogenase substrate *in vitro*^36^. However, the Fe protein from *Klebsiella pneumonia* can partially complement the function of ChlL in *Chlamydomonas reinhardtii*^38^. Structures of COR and F430 have yet to be solved, and few biochemical studies have been conducted on these enzymes^30^.

The light-driven and nuclear-encoded LPOR is a single-polypeptide enzyme, unlike the light-independent multi-subunit DPOR. LPOR and DPOR share no homologous amino-acid sequences, while both catalyze the reduction of the same substrate despite *via* different reaction mechanisms^39^. Intriguingly, LPOR and DPOR coexist in many oxygenic photosynthetic organisms such as *Chlamydomonas reinhardtii* and *Pinus mugo*, but not angiosperms (e.g., *Arabidopsis thaliana*) where DPOR is absent^40^. This may imply that their coexistence during evolution was required to guarantee sufficient chlorophyll production under varying light and oxygen conditions^40^. In contrast, LPOR is widely distributed among aerobic anoxygenic phototrophic bacteria, angiosperms, and non-photosynthetic organisms due to possible horizontal gene transfer^40^. Oxygen-sensitive DPOR probably originated prior to LPOR in an anaerobic environment^37,39^. However, the lack of similarity in primary amino-acid sequences of DPOR and LPOR suggests that they are evolutionarily unrelated. This perception contradicts their functional evolution, as LPOR has evolved to compensate for, complement, and/or replace DPOR for photosynthesis in cyanobacteria. On the other hand, DPOR resembles nitrogenase in primary amino-acid sequence and structures. However, whether LPOR has evolved from photosynthesis-unrelated nitrogen-fixation nitrogenase enzymes and nitrogenase-like proteins including DPOR remains unknown.

In this study, using structural alignment tools, we uncover the conserved 3D-structural domains with distinctive primary amino-acid sequences and a previously undescribed evolutionary relationship between LPOR and nitrogenase-like proteins. We demonstrate that an archaic one-subdomain protein precursor appears to be the common ancestor for nitrogenase-like protein complexes DPOR, COR and the chlorophyll biosynthesis key enzyme LPOR. Moreover, we envisage that the innovative structure-guided evolutionary analysis approach, designated “3D-structural similarity implying evolutionary relationship” strategy, may be exploited to design novel functional proteins for engineering high-efficient BNF, photosynthesis, and beyond.

## RESULTS

### Cyanobacterial LPOR and photosynthetic bacterial DPOR share structural similarities

Eukaryotic LPORs have evolved from cyanobacterial LPORs and DPORs, and DPORs likely originated in anoxygenic photosynthetic bacteria, such as purple and/or green sulfur bacteria^31,41^. On the other hand, Cyanobacteria are thought to evolve from non-photosynthetic *Sericytochromatia* into which photosynthetic reaction centers were horizontally transferred^42^. Taking these into account, we proposed that DPOR may be directly acquired through horizontal transfer, and LPOR emerged later through evolution in cyanobacteria. Given the similar enzymatic activities of the single-polypeptide LPOR and the octameric DPORs in reducing protochlorophyllide to chlorophyllide for (bacterio)chlorophyll biosynthesis, we hypothesized that DPOR and LPOR might share structural similarities although they lack primary amino-acid sequence identity. To test this hypothesis, we used AlphaFold2 to predict the non-modeled regions of two experimentally crystalized cyanobacterial LPORs PDB: 6R48 and PDB: 6L1H, and obtained the full LPOR structure (Figure 1A). Using a similar approach, we optimized the conserved structure of DPORs from the photosynthetic purple bacteria, i.e., BchL from *Rhodobacter sphaeroides* and BchN/BchB from *R. capsulatus*, (Figure 1B). The Rossmann folds^8,43–45^ are evident in our improved LPOR and DPOR structures (Figure 1A, 1B, S1A, and S1B). To uncover whether LPOR and DPOR share structural similarity, we aligned structurally the cyanobacterial LPOR (Figure 1A) and the three individual subunits BchL, BchN and BchB of the *Rhodobacter* DPOR complex (Figure 1B). To our surprise, BchL was found to share high structural similarities with LPOR, having a DALI Z-score of 5.5, although slightly lower in case of BchN vs LPOR and BchB vs LPOR, having a DALI Z-score of 2.6 or 4.9, respectively (Figure 1C). However, all these DALI Z-scores are markedly higher than the DALI Z-score of 2 for similarity threshold, indicating the bona fide existence of related structures in LPOR vs different subunits of DPOR.

**Figure 1.**
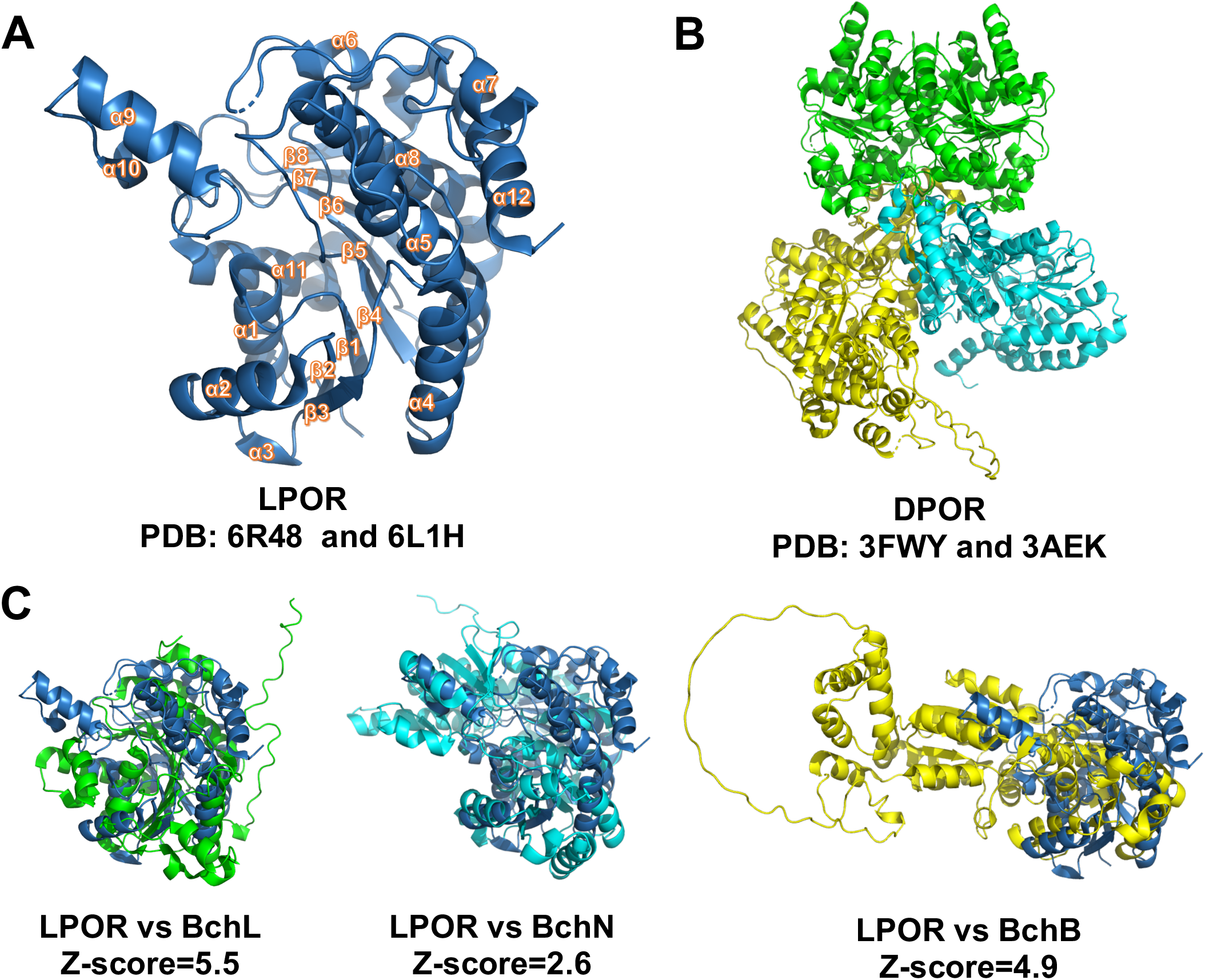
LPOR and DPOR share structural similarities. (A) Ribbon diagrams showing the optimized 3D structure of LPOR. Twelve α-helices and eight β-sheets were generated by integrating crystalized structures of two cyanobacterial LPORs (PDB: 6R48 + 6L1H) with the assistance of AlphaFold2 predictions to supplement the non-crystalized regions in the LPOR proteins. (B) Ribbon diagrams showing the optimized 3D structure of DPOR consisting of three subunits BchL (green), BchN (cyan), and BchB (yellow) from the photosynthetic purple bacterium i.e., BchL from *Rhodobacter sphaeroides* (PDB: 3FWY) and BchN/BchB from *R. capsulatus* (PDB: 3AEK). (C) Structural alignments of LPOR with the three subunits BchL, BchN, and BchB of DPOR in ribbon representation using US-align, respectively. LPOR is shown in blue, BchL in green, BchN in cyan, and BchB in yellow. The structural similarity was conducted using the DALI server, and the structural similarity threshold is a DALI Z-score of 2.

### LPOR, nitrogenase, COR and F430 possess similar protein folds

The three subunits NifH, NifD, and NifK of Nitrogenases possess similar amino-acid sequences with the three subunits of two nitrogenase-like proteins, i.e., BchL, BchN and BchB of DPOR, and BchX, BchY and BchZ of COR, respectively^46–48^. The two subunits CfbC and CfbD of F430 also show sequence similarity to BchL and BchN of DPOR^46,47^. These imply that DPOR may share structural similarities with nitrogenase, COR and F430. As nitrogenase isoenzymes exhibit a high degree of structural conservation across different organisms, the Fe-only (Anf) nitrogenase from the anoxygenic photosynthetic *R. capsulatus* was selected for analysis in this study due to its being the only nitrogenase whose crystal structure solved from phototrophic bacteria (Figure 2A)^49^. To determine their structural similarities, we first predicted the structures of multimeric COR and F430 using AlphaFold2 multimer. Prediction accuracy was measured using the combination of ipTM (interface predicted template modeling) and pTM (predicted template modeling) scores. The AlphaFold2-predicted structures of multimeric COR and F430 with the highest confidence (a score of 0.78 and 0.9, respectively) were selected for further analysis (Figure 2B, 2C). Next, we conducted pairwise comparisons of structures of all subunits of DPOR, nitrogenase, COR and F430 using the DALI server. As shown in Figure 2D, these subunits were structurally clustered into component 1 and component 2 in the DALI fold dendrogram. Component 2 includes BchL, AnfH, BchX, and CfbC which correspond to the ATPase domain, while component 1 contains BchN, BchB, AnfD, AnfK, BchY, BchZ, and CfbD. Intriguingly, CfbD was outbranched from other members of component 1, close to the center of the phylogenetic tree (Figure 2D), in line with the previous suggestion that F430 might be an ancestor for nitrogenase and other nitrogenase-like enzymes^46^. Excluding CfbD, component 1 includes both α (AnfD/BchN/BchY) and β (AnfK/BchB/BchZ) subunits, suggesting that these subunits may have originated from a common ancestor after a duplication event, supported by previous findings (Figure 2D)^46,50^.

**Figure 2.**
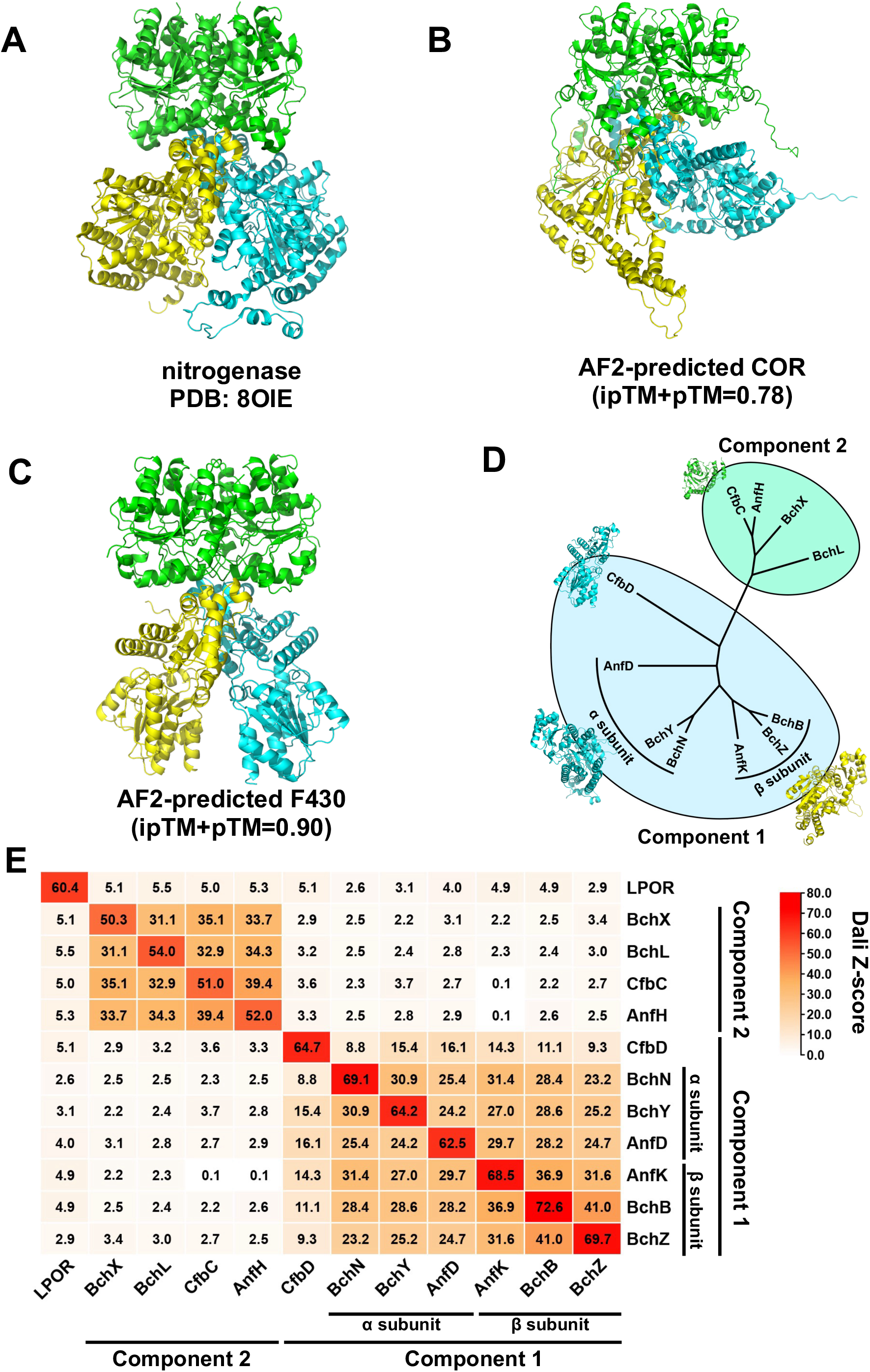
LPOR and different subunits of nitrogenase, COR and F430. (**nitrogenase-like proteins) share structural similarities** (A) Ribbon diagrams showing the 3D structure of the iron-only nitrogenase from the phototrophic bacterium *Rhodobacter capsulatus.* The structure of the iron-only nitrogenase was generated based on PDB identifier 8OIE with the assistance of AlphaFold2 predictions. The subunit AnfH is shown in green, AnfD in cyan and AnfK in yellow. (B) Ribbon diagrams showing the AlphaFold2-predicted structure of COR with the highest ipTM+pTM score of 0.78. The structure of COR was predicted using AlphaFold2 multimer. The subunit BchX is shown in green, BchY in cyan, and BchZ in yellow. (C) Ribbon diagrams showing the AlphaFold-predicted structure of F430 with the highest ipTM+pTM score of 0.9. The structure of F430 was predicted using AlphaFold2 multimer. The first subunit CfbC is shown in green, CfbD in cyan, and the second CfbD subunit in yellow. (D) Hierarchical cluster analysis showing the DALI fold dendrogram among the subunits of DPOR, nitrogenase, COR, and F430. (E) Heatmap showing the structural similarity between LPOR and different subunits of nitrogenase, COR and F430 using the DALI server. The structural similarity was evaluated using DALI Z-scores. A threshold DALI Z-score of 2 was applied to determine the structural similarity.

Furthermore, all subunits in component 1 exhibit structural similarities (DALI Z-scores 8.8-41.0), and subunits except CfbD in component 1 have high similarity (DALI Z-scores 23.2-41.0) for all comparisons (Figure 2E). Compared to the structural similarity (DALI Z-scores, 24.2-30.9) among α subunits in component 1, the β subunits exhibit a higher degree of structural similarity (DALI Z-scores 31.6-41.0). These data suggest that the β subunits might have undergone strong selection pressures during the evolution of different nitrogenase-like proteins, resulting in their relatively more conservation. In contrast, the α subunits, which are involved in substrate binding, have experienced divergent selection pressures to adapt to different substrates, leading to increased structural specificity (Figure 2E). On the other hand, among α subunits, BchN and BchY exhibit the highest structural similarity with a DALI Z-score of 30.9. The highest structural similarity among β subunits is between BchB and BchZ, with a DALI Z-score of 41.0, indicating that COR and DPOR share highly similar structural folds. This is consistent with the fact that BchB and BchZ have the highest amino-acid sequence identity and similarity of 26.10% and 33.78%. In addition, in component 2 the structural similarities between the subunits are also high, with DALI Z-score exceeding 31 for all pairwise comparisons (Figure 2E). Interestingly, we observed that LPOR and other enzymes display structural similarity with DALI Z-scores ranging from 2.6 to 5.5, and individual subunits within component 2 have DALI Z-scores greater than 2 when compared to any subunit in component 1 (Figure 2E). The three subunits BchL exhibit the highest structural similarity to LPOR with a DALI Z-score of 5.5 (Figures 1D and 2E). Collectively, these findings indicate that functionally divergent LPOR and nitrogenases, DPOR, COR, and F430 have evolved similar protein folds, implying that structurally they may share a distant common ancestor.

### All subunits of nitrogenase-like proteins have evolved from the single ancient CfbC subunit

A structural subdomain with a similar Rossmann fold has previously been identified both in component 1 of DPOR and nitrogenase^20,43,51^. Interestingly, a similar structural fold with α-helices and β-sheets was observed in LPOR, an SDR protein with the typical Rossmann fold (Figure 1A). Thus, we dubbed this structural fold in LPOR, DPOR and nitrogenase as the Core (β-sheet)-Guard (α-helix) subdomain (Figures 1A and S1A). We noticed that the Core-Guard subdomain was present in all subunits of nitrogenase-like protein (Figures 1B, 2A-2C, S1B and S2A-S2C). Interestingly, all subunits in component 2 comprise only a single Core-Guard subdomain, while those in component 1 contain three except CfbD has two such subdomains (Figures 1A-1C, 2A-2C, S1A, S1B and S2A-S2C).

Given that all subunits of component 1 contain at least two Core-Guard subdomains, we hypothesized that evolutionary relationships exist among these subdomains. To test this assumption, we first split the subunits of component 1 into individual subdomains. CfbD, referred to as a two-subdomain subunit, was divided into CfbD_N and CfD_C representing the N- and C-terminal portions of the protein (Figure S3A). The remaining subunits in component 1, designated three-subdomain subunits, were divided into N-terminal, Middle (M) and C-terminal subdomains (Figure S3B-S3D). Pairwise structural similarities among subdomains of component 2 and component 1 subunits were analysed using the DALI server. As shown in Figure 3A, these subdomains were categorized into four clusters. Specifically, all subdomains in Cluster 1 correspond to component 2 subunits, while Clusters 2, 3, and 4 contain the subunits of component 1 (Figure 3A). Remarkably, the N- and C-terminal (except CfbD_C), and M subdomains of component 1 formed three independent clusters (Figure 3A), suggesting that each individual subdomain in the same cluster has originated from a common three-subdomain ancestor, dubbed three-subdomain precursor hereafter. Cluster 1, positioned distinctly apart from the other three clusters, is regarded as the outgroup, implying that all subdomains may have originated from Cluster 1 (Figure 3A).

**Figure 3.**
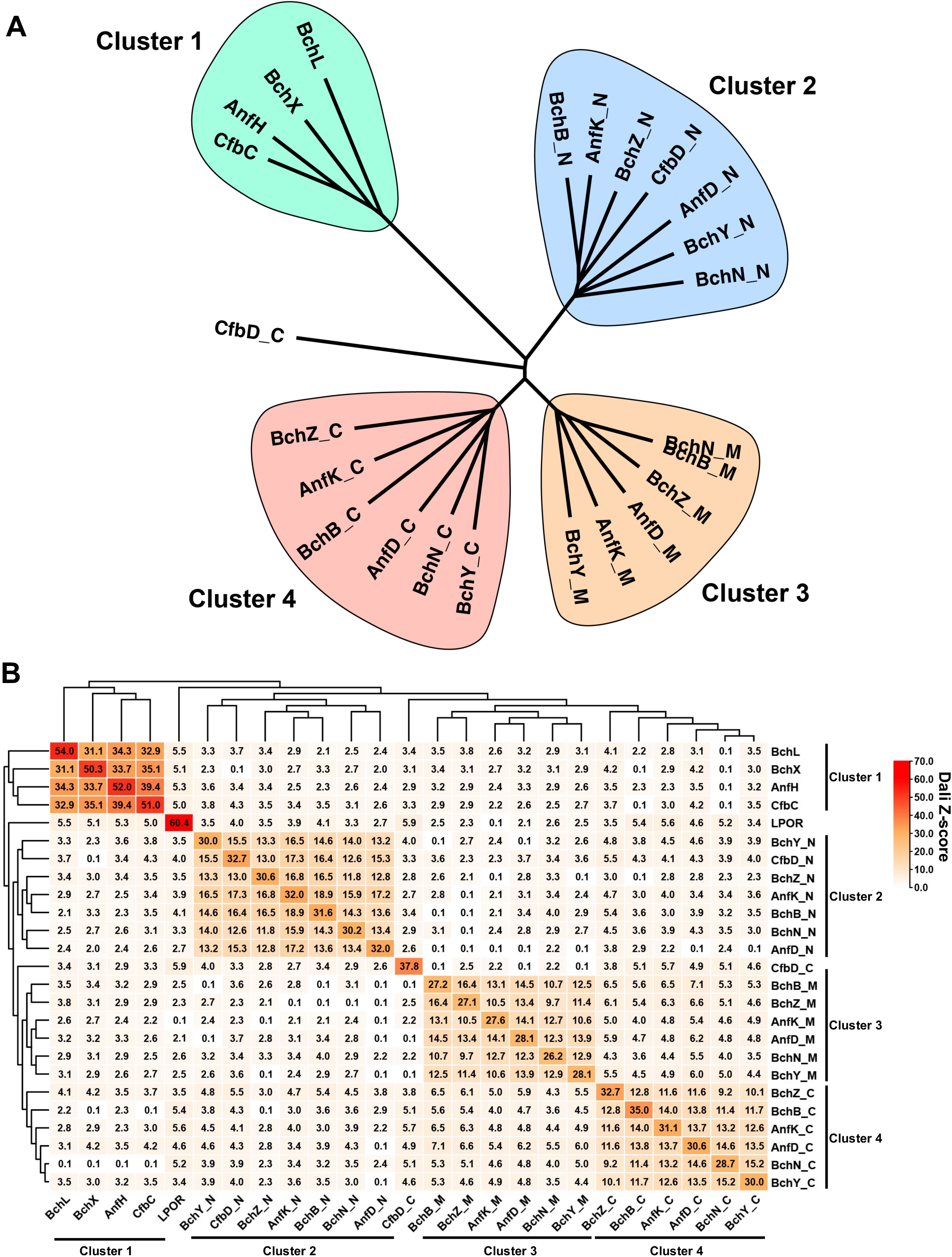
Structural similarities among subdomains of component 1 and component 2 subunits of nitrogenase-like proteins and LPOR. (A) Hierarchical cluster analysis showing the DALI fold dendrogram among different subdomains of subunits of DPOR, nitrogenase, COR, and F430 and LPOR. The CfbD subunit was split into N-terminal (CfbD_N) and C-terminal subdomain (CfbD_C) based on its structure, and subunits BchN, BchB, BchY, BchZ, AnfD, and AnfK of component 1 were structurally divided into the N-, Middle and C-terminal subdomains. Structural similarities among these subdomains were identified by the DALI server. (B) Heatmap showing the structural similarities among the different subdomains of subunits of DPOR, nitrogenase, COR, and F430 and LPOR using the DALI server. The structural similarity was evaluated and determined by a threshold DALI Z-score of 2.

Although CfbD_C is absent in Cluster 4, it has structural similarity with the C-terminal subdomains of three-subdomain subunits in Cluster 4 (DALI Z-scores 3.8-5.7), but bears no or limited similarity with the M subdomains of three-subdomain subunits in Cluster 3 (DALI Z-scores 0.1-2.5) (Figure 3A and 3B), suggesting that the C-terminal region of all three-subdomain subunits in component 1 and CfbD_C share a structurally related common ancestor. The M and C-terminal subdomains of three-subdomain subunits exhibit high structural similarity (DALI Z-scores 3.5-7.1) (Figure 3A and 3B), implying that the M and C-terminal subdomains of the three-subdomain subunits have likely originated from a common two-subdomain ancestor, dubbed two-subdomain precursor hereafter, resulted from possible genetic duplication. We also noticed some structural similarity between CfbD_N vs CfbC (DALI Z-score 4.3), and CfbC vs CfbD_C (DALI Z-score 3.3) (Figure 3B). Using CfbD_N as an input to search against the Swiss-Prot database specific to *Methanosarcina acetivorans*, an anaerobic methanogenic archaeon, by Foldseek^52^, we found that CfbC in Cluster 1 or component 2 is the fourth-highly ranked output (Figure S4). These findings indicate that the three-subdomain subunits in component 1 are evolutionarily related to all subunits in component 2, and CfbC is evolutionarily close to their common ancestor. We speculate that the two-subdomain precursor may have evolved from a single-domain ancestor, so called one-subdomain precursor, *via* a genetic duplication event. To test this idea, we used US-align and compared the structures between CfbC and CfbD_N/_C which were supposed to evolve from the one- or two-subdomain precursor, respectively. We found that most secondary structures in CfbD correspond to those in CfbC (Figure 4A). We also compared the AlphaFold2-predicted structures of CfbD_N/CfbD_C with CfbC, and uncovered CfbD_N and CfbD_C share structural similarities with CfbC (DALI Z-scores 4.3 and 3.3, respectively) (Figure 4B). We further generated the simulated models of CfbD_N and CfbD_C by assembling the amino acid sequence of CfbC, which exhibits structural similarity with CfbD. The AlphaFold2-predicted structures of the simulated CfbD_N and CfbD_C display similarity to the AlphaFold2-predicted CfbD structure (DALI Z-scores 5.6 and 4.7) (Figure S5). These findings suggest that the two-subdomain precursor as the ancestor of CfbD has evolved from the CfbC ancestor of the one-subdomain precursor through self-genetic duplication.

**Figure 4.**
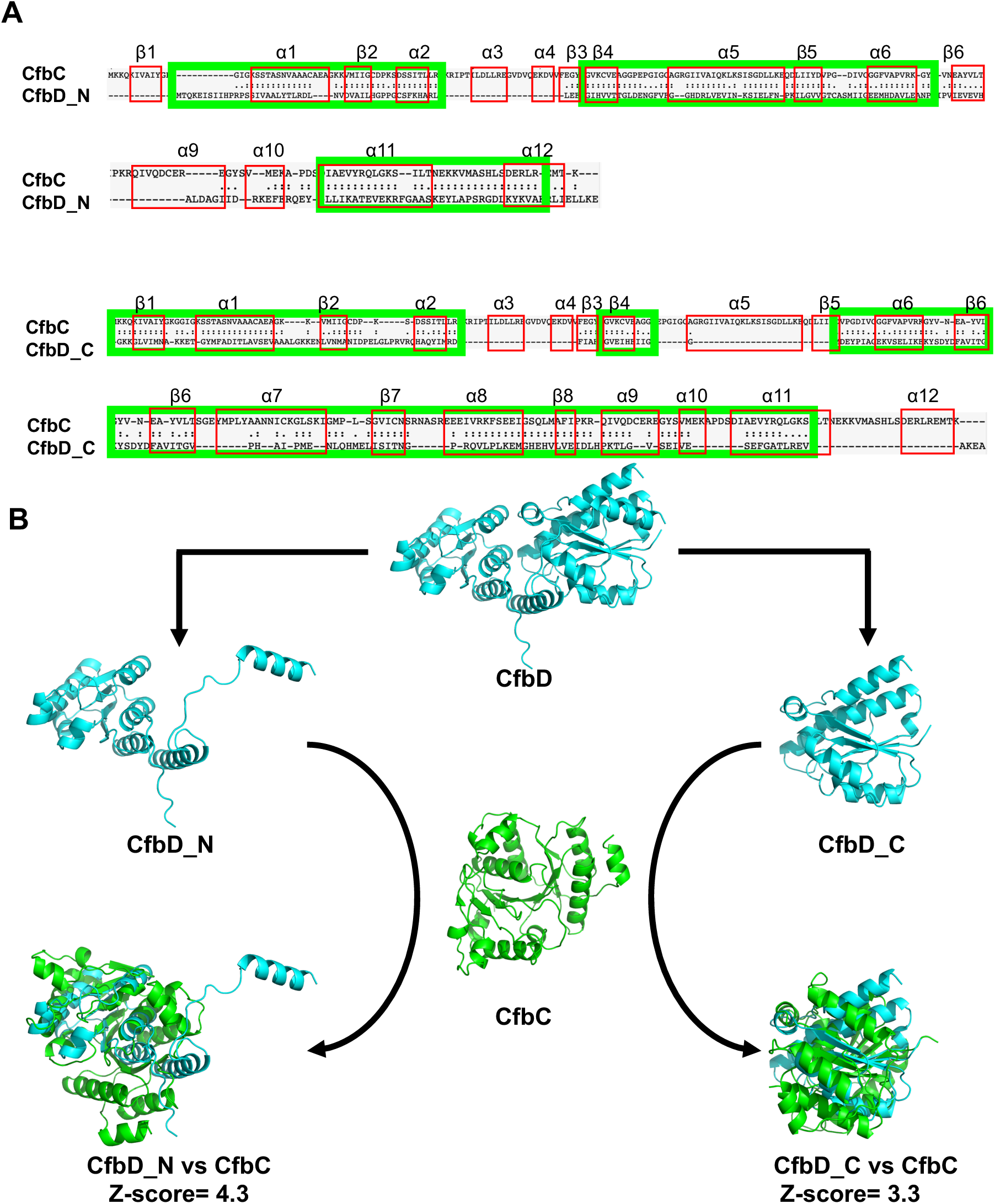
CfbD_N and CfbD_C share 3D-structural similarities with CfbC. (A) US-align based structure alignments of CfbC vs CfbD_N (Top) and CfbC vs CfbD_C (Bottom). Red box indicates the secondary structure of CfbC. Green box indicates regions with high 3D-structural similarities. The alignments highlight amino acid fragments showing significant structural similarities. “:” denotes residue pairs of d < 5.0 Angstrom, “.” denotes other aligned residues. (B) Comparison diagram of the AlphaFold2-predicted CfbD_N (Top left) and CfbD_C (Top right) structures with AlphaFold2-predicted CfbC (Middle) structure generated by US-align. US-align based superpositions of CfbD_N vs CfbC (Bottom left) and CfbD_C vs CfbC (Bottom right) are shown in ribbon representation. The structure of CfbC is in green, and the structures of CfbD_N and CfbD_C are in blue. The structural similarity threshold is a DALI Z-score of 2 generated by DALI server.

### Evolution of LPOR from ancient multiple subunit COR

Structures of the three subdomains with Rossmann folds in NifD and NifK were phylogenetically related^47^. This has prompted to postulate that these subdomains have originated from an ancient single-domain protein through genetic duplication, a process crucial for the evolution of complex nitrogen-fixing systems^51^. With the successful characterization of analogous structures among different subdomains of nitrogenase-like proteins and LPOR, we were able to deduce how a primordial single-subdomain precursor has evolved into multiple-subunit nitrogenase-like proteins and LPOR which share no identical primary amino-acid sequences. As shown in Figure 5 and Video S1, the one-subdomain precursor is the ancestor of all subdomains, and the two-subdomain precursor is formed through potential protein duplication^50^ and recombination^53,54^ events. Subsequently, the entire component 1 originates from a common two-subdomain precursor. One branch of the two-subdomain precursor evolves into CfbD without undergoing the C-terminal subdomain duplication, while the other branch duplicates the C-terminal subdomain to form a three-subdomain precursor. The α and β subunits have diverged from the three-subdomain precursor and become the subunits of nitrogenase, COR, and DPOR of component 1 to meet the requirements of biological function differentiation. Following an extra subdomain duplication, the M subdomain endures further diversification, leading to low structural similarity with CfbD, but retaining the structural similarity with the C-terminal subdomain. The emergence of the M subdomain signifies an adaptation to increased functional complexity, possibly related to survival in new environments.

**Figure 5.**
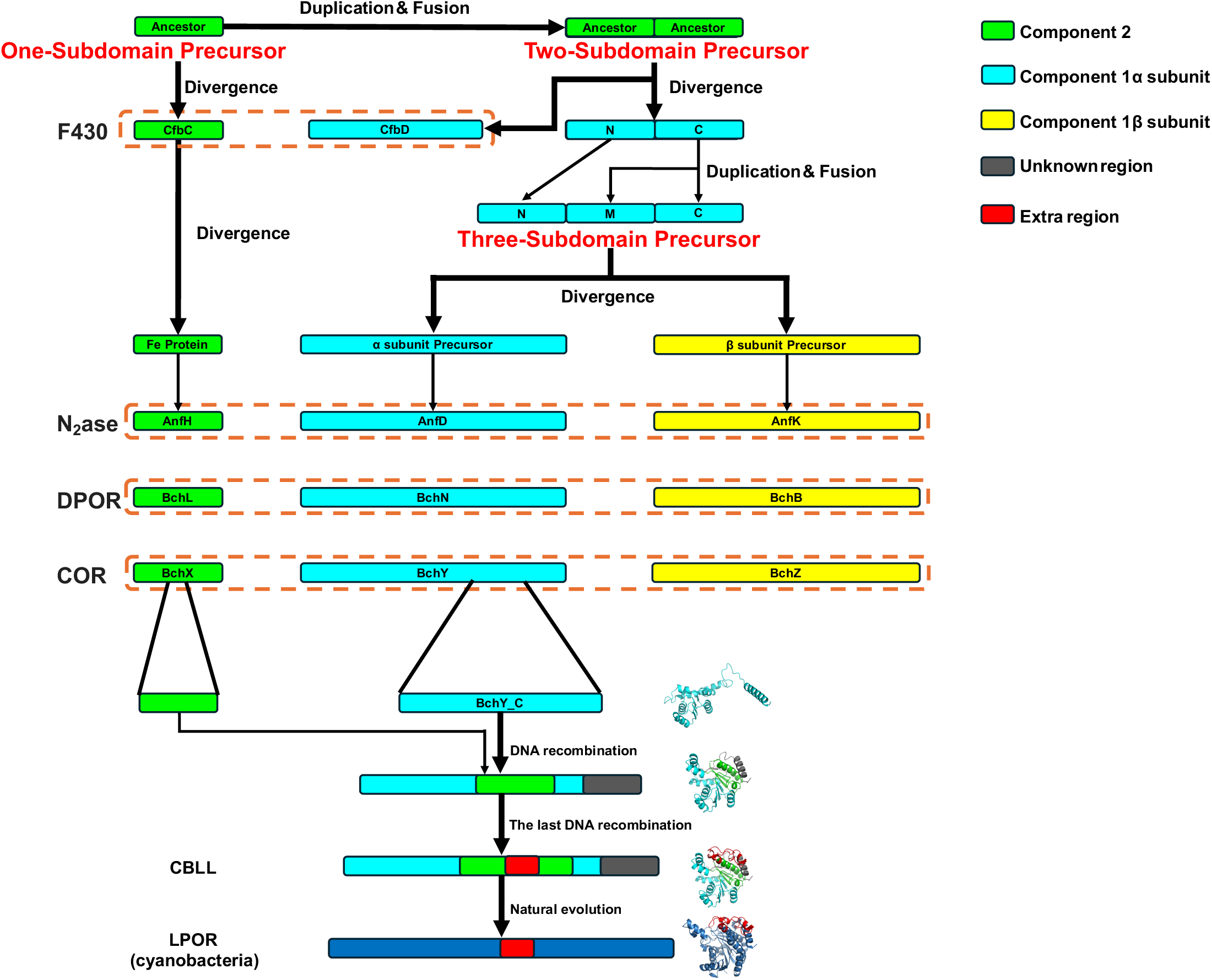
Schematic representation of the deduced evolutionary path of LPOR and nitrogenase-like proteins. The evolutionary trajectory of LPOR originates from ancient nitrogenase-like proteins, involving key genetic recombination and duplication events. The path starts with a hypothesized one-subdomain precursor, which evolved through gene duplication and recombination to form various nitrogenase-like protein complexes, such as DPOR and COR. A two-subdomain precursor emerged from the one-subdomain ancestor and evolved into a three-subdomain precursor, leading to the divergence of multi-subunit nitrogenase-like proteins. COR evolved into LPOR through twice genetic recombination. This schematic representation provides an overview of the evolutionary relationships among nitrogenase-like proteins and the structural innovations that led to the development of LPOR in photosynthesis.

Structural comparisons between *Escherichia coli* 7-α-hydroxysteroid dehydrogenase (AHI) and LPOR revealed a shared conformation fold across the entire sequence, except for one region called “extra loop”. This “extra loop” corresponds to amino acids 217-252 in PORA and 225-260 in PORB^55^, implying that this region is likely inserted into the ancestor of LPOR during evolution. Importantly, another extended region of approx. 47 amino-acids (including α6-α7 and part of α8) that is slightly longer than the “extra loop” identified in PORA and PORB, is absent in nitrogenase and all subunits of nitrogenase-like proteins, dubbed “extra region” (Figure 5 and S6A). To define the evolutionary history of LPOR, we superimposed the entire LPOR structure onto each subunit of nitrogenase-like proteins and onto each Core-Guard Subdomain of each subunit (Figure S6A). Surprisingly, except for the extra region and the ∼45 amino-acid C-terminal region of LPOR, nearly all structures match their counterparts in nitrogenase-like proteins, implying that approx. 71% (92/318 amino-acids) structure of the 318 amino-acid LPOR likely originate from nitrogenase-like proteins (Table 1). To trace the origin of LPOR, we established four principles (1) structural continuity, (2) similarity score, (3) amino acid substitution ratio, and (4) evidence supporting the presence of extra regions. These principles are guided by the structural similarity observed in the Core-Guard subdomains across different subunits. Following these rules, we generated four ancestral-like LPOR sequences by employing sequences from four types of nitrogenase-like proteins (1) DPOR-based LPOR-like protein (DBLL), (2) COR-based LPOR-like protein (CBLL), (3) nitrogenase-based LPOR-like protein (NBLL), and (4) F430-based LPOR-like protein (FBLL) (Figure S6B). DBLL, CBLL, NBLL, and FBLL represent 70.75%, 71.01%, 67.92%, and 60.38% of the original LPOR sequence, respectively. The remaining regions, where structural similarity could not be traced, retain the original LPOR sequence. Subsequently, using AlphaFold2 we generated and then compared the structures of DBLL, CBLL, NBLL, and FBLL with the actual structure of LPOR using the DALI server (Figure S6B and S6C). We used the afore-described four principles and evaluated the AF2-predicted structures of DBLL, CBLL, NBLL, and FBLL for tracing the evolution path of the naturally occurring LPOR. Based on these comparative structural analyses, the artificial CBLL was found to be the optimal ancestral-like LPOR, evidenced by having the best continuity (2 segments), the highest amino acid substitution ratio (71.01%), a high similarity score (20.1), and the insertion of the extra region (Table 1 and Figure S6A-S6C). The data suggest that LPOR has naturally originated from COR.

**Table 1.** Evaluation for four ancestral-like LPORs based on structural similarity.

| Evaluation Criteria | DBLL | CBLL | NBLL | FBLL |
| --- | --- | --- | --- | --- |
| Sequence Continuity | 5 segments | 2 segments | 4 segments | 4 segments |
| Similarity Score (from DALI server) | 21.2 | 20.1 | 12.0 | 20.0 |
| Amino Acid Substitution Ratio | 70.75% | 71.01% | 67.92% | 60.38% |
| Evidence Supporting Extra Region | NO | YES | NO | NO |

To confirm this, we carried out detailed structural alignments using US-align and found that β1-α1-β2-α2-α3-β3-α4-β4 of LPOR corresponds to β1-α1-β2-α2-β3-α3-β4 in the BchY_C subdomain of COR (Figure S6A). LPOR α5-β5 and α8-β6 align with α7-β6 or α8-β7 in the BchX of COR, respectively (Figure S6A). The extra region (α6-α7) is absent in the BchX of COR, but accommodation of α6-α7 is essential for the evolution of LPOR^10,55^. In addition, we observed that α9-α10 and α11 correspond to α4-β5-α5-α6 of BchY_C of COR, and there is no similar structure in COR corresponding to β7-β8-α12 of LPOR (Figure S6A). Using the organization and order information of structural polypeptide derived from COR in CBLL, we deduced the stepwise evolution of LPOR from COR, in which *BchY_C* fragment as ancestral part undergoes two rounds of genetic recombination with *BchX* fragement, and an insertion event in *BchX* fragement in the final step under natural selection to evolve the present LPOR (Figure 5).

**Video S1.**
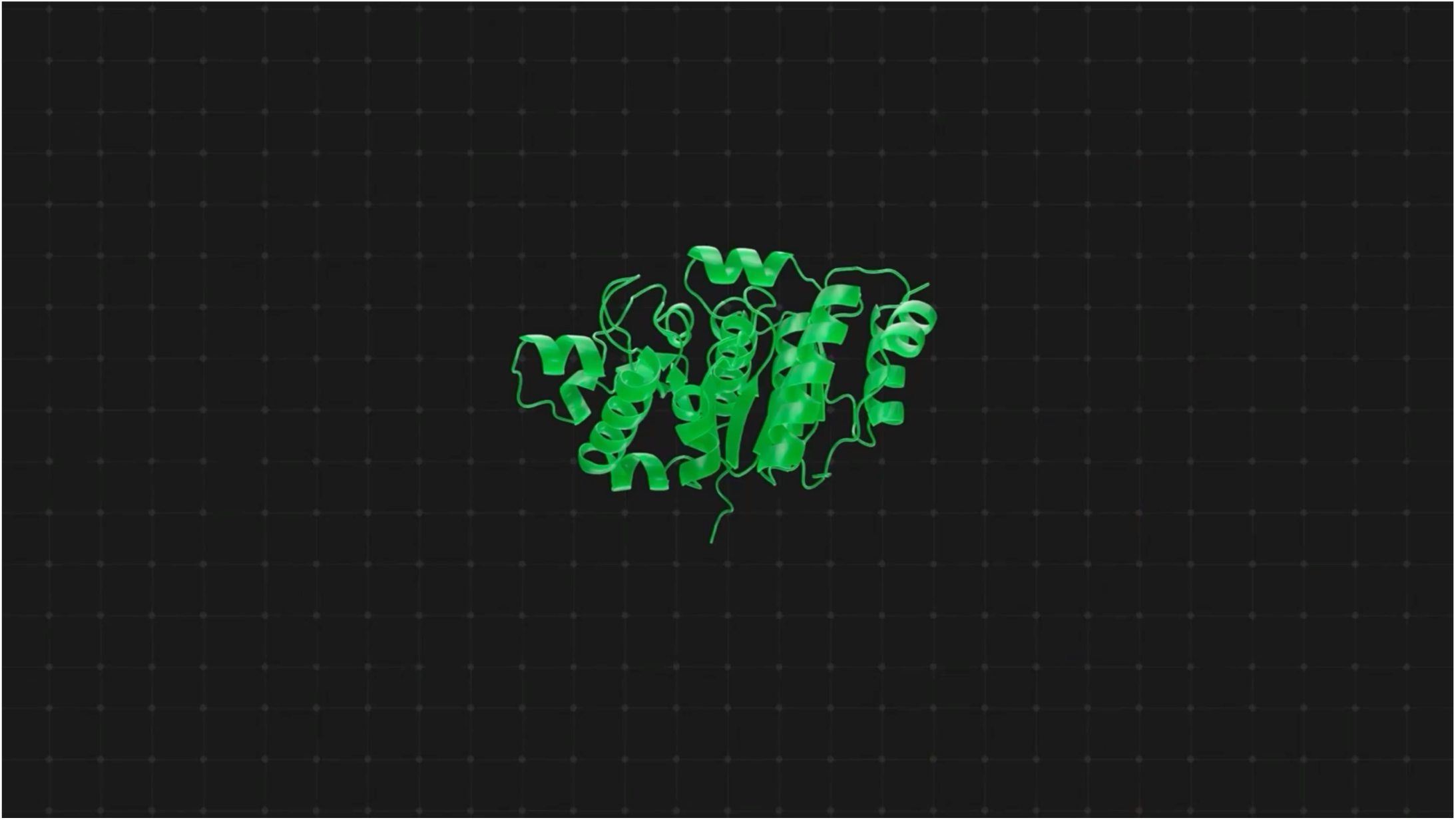
Structural origins and evolution of nitrogenase-like proteins, related to Figure 5.

**Video S2.**
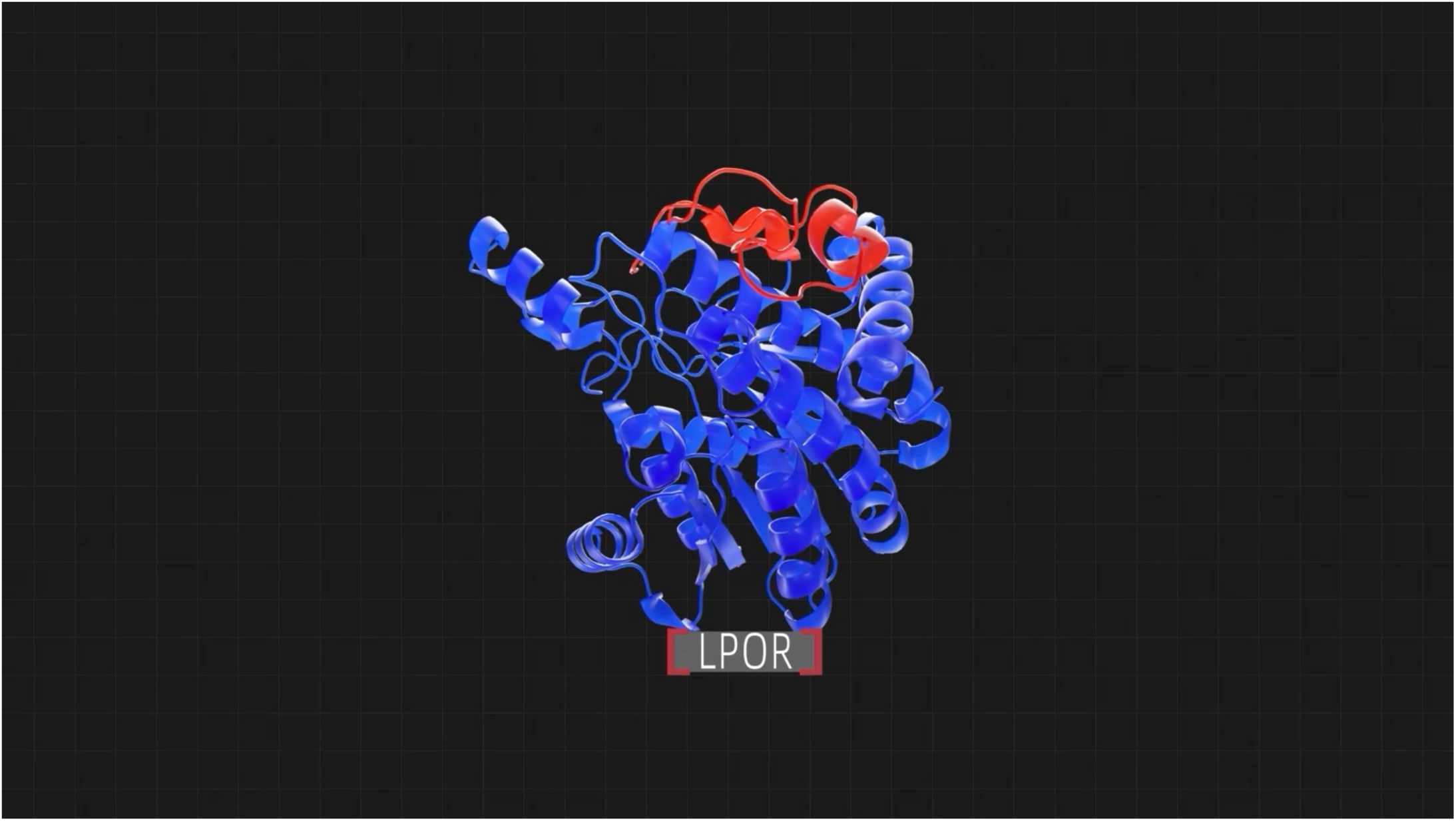
Structural origins and evolution of LPOR, related to Figure 5.

### Animal retinol dehydrogenase is an unexpected intermediate in the COR-to-LPOR evolution path

As COR evolved into LPOR potentially through two genetic recombination events, we hypothesized that an intermediary protein between COR and LPOR should exist in organisms lacking both COR and LPOR (Figures 5, 6A, and Video S2). To test this conception, we aligned the structure of LPOR against the AlphaFold Swiss-Prot database using Foldseek^52^. Intriguingly, one of the top matches is human retinol dehydrogenase (RDH) 13 which shares high structural similarity (TM-score of 0.792) and 31.2% amino-acid sequence identity (Figure S7). We designated the AlphaFold2-predicted intermediate protein, which lacks the extra region, as COR-based RDH-like protein (CBRL) (Figures 5, 6A, and Video S2). We further performed a BLASTP search of LPOR against the NCBI NR database and found that in addition to the human RDH, the RDH (Accession No. XP_019625646.1) from lancelet (*Branchiostoma belcher*) exhibits 32.41% amino-acid identity with LPOR over 97% sequence coverage at an E-value of 1e-38. Such high amino-acid sequence identities between LPOR vs lancelet/human RDHs suggest that the LPOR-related sequences were preserved in animals before the divergence of invertebrates (e.g. lancelet) and vertebrates (e.g. humans). Animal RDHs are closely related to LPOR and have further evolved within the animal kingdom. Significantly, animal RDHs lack the extra region present in LPOR (Figure 5), yet they still retain more than 30% amino-acid sequence identity with plant LPORs despite hundreds of millions of years of divergent evolution. It is worthwhile noting that *B. belcheri* is a primitive chordate, and a model organism for studying early animal development and evolutionary biology. Lancelet also lacks a vertebral column yet displays vertebrate traits and occupies a key evolutionary niche between invertebrates and vertebrates, making it an ideal subject for studying transitional phases^56,57^. Taken together, our data demonstrate that RDHs have functioned as an essential intermediate for COR to evolve into LPOR, as well as highlight extraordinary cross-kingdom conservation among these enzymes during COR-to-LPOR evolution (Figures 5 and 6A).

**Figure 6.**
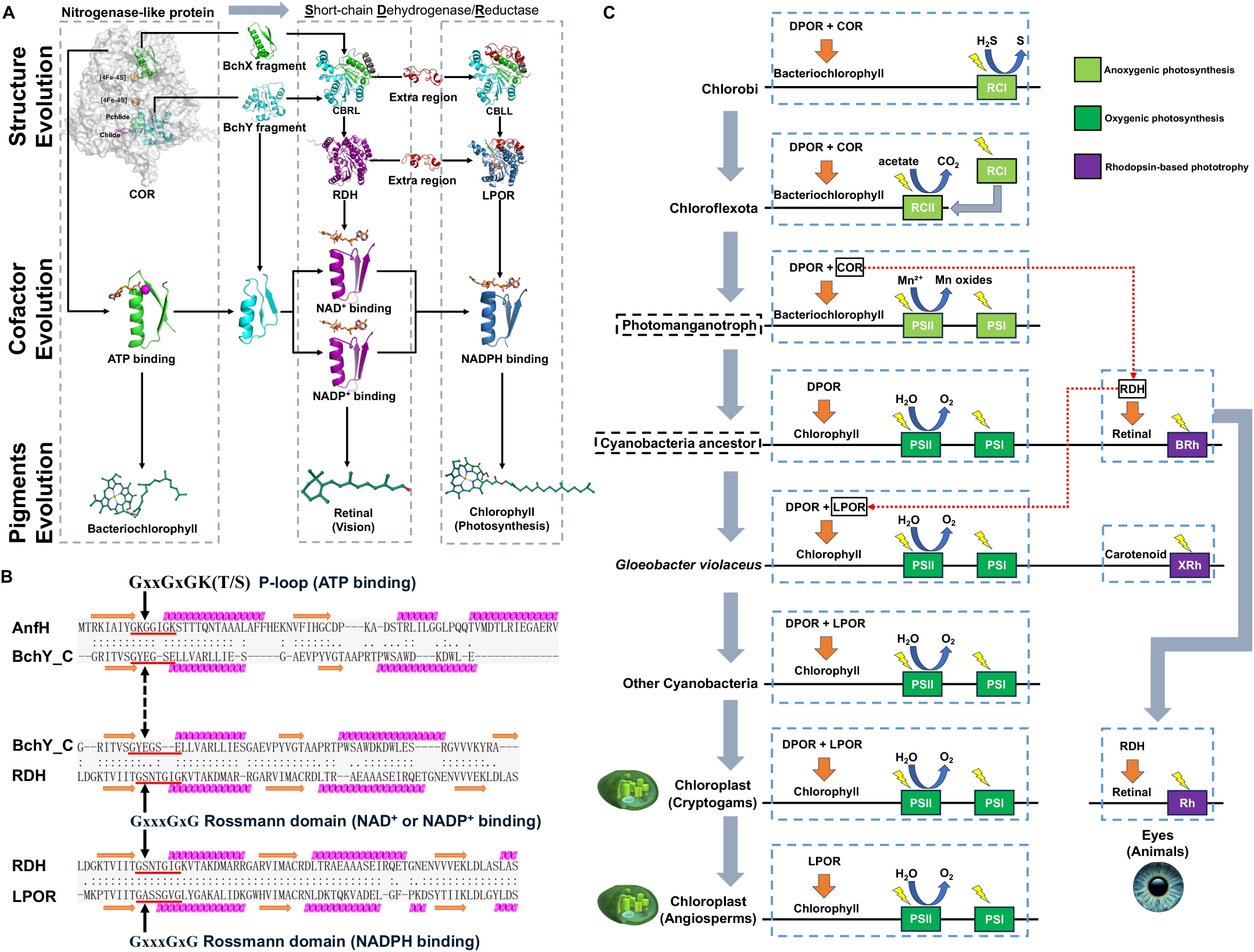
Schematic representation of the deduced evolutionary pathways of energy-converting pigments and associated enzymes. (A) Schematic representation of the evolutionary transition from COR to LPOR *via* RDH. Insertion of the extra region (red) into the simulated RDH led to formation of a hypothetical reductase. Two structural segments of COR contributing to the generation of COR-derived RDH-like LPOR are highlighted green and blue. The positions of two small molecules, Chlide (pink) and Pchlide (green), are depicted. The Chlide location is derived from the molecular docking results with the best docking mode exhibiting an affinity of - 11.5 kcal/mol. Pchlide is spatially positioned in COR based on the structural similarity between COR and DPOR. AlphaFold2-predicted structure of RDH from *Branchiostoma belcheri* is shown purple. The docking of NAD^+^/NADP^+^ cofactors into RDH was displayed based on AlphaFold3-predicted structural models. The COR-to-LPOR evolution also involves cofactor changes and binding structure adjustment, specifically transitioning from ATP binding in component 2 (AnfH as a representative example) to NAD^+^/NADP^+^ binding in RDH and NADPH binding in LPOR. Pigment evolution also occurs, with a shift from bacteriochlorophyll to retinal and chlorophyll, reflecting adaptations for diverse biochemical roles. (B) US-align based structure alignments of component 2 (AnfH as a representative) ATP-binding P-loop (GxxGxGK(T/S)), N-terminal of BchY_C subdomain and the NAD^+^/NADP^+^/NADPH binding GxxxGxG Rossmann domain. The alignments highlight amino acid fragments showing significant structural similarities. “:” denotes residue pairs of d < 5.0 Angstrom, “.” denotes other aligned residues. (C) The evolution of phototrophic metabolism from anoxygenic photosynthesis (light green) to oxygenic photosynthesis (dark green) and rhodopsin-based phototrophy (purple).

### Cofactor evolution in nitrogenase-like proteins and RDH/LPOR

To gain a deep insight into the COR-to-LPOR evolution, we analyzed and unraveled the structural segments of COR that contribute to the evolution of CBLL (Figure 6A and Video S2). Intriguingly, the segment from BchX is positioned closely to the [4Fe-4S] cluster (mapped from the crystal structure of DPOR), while the segment from BchY is located proximately to the ligand Chlide (from molecular docking) or Pchlide (mapped from the structure of DPOR) (Figure 6A and Video S2). These findings indicate that all of the functional RDH and LPOR structures originate from the key COR sequences. Considering nitrogenase-like proteins bind to ATP, whereas RDH to NAD^+^ or NADP^+^, and LPOR to NADPH to catalyze, such structural correspondence suggests that the biochemical process about how light energy catalysis is invoked by LPOR is likely to have evolved after the ATP-utilizing catalytic mechanism.

The NADPH- and NAD^+^/NADP^+^-binding motifs in LPOR and RDH contain a GxxxGxG Rossmann domain which originates from the N-terminal of BchY_C (Figure 6A, 6B and S6A). BchY_C and all other subunits of nitrogenase-like proteins can be traced back to the ancestral one-subdomain precursor. Thus, the N-terminus of BchY_C is probably derived from the N-termini of component 2 subunits, which include the ATP-binding GxxxGxG Rossmann domain and the P-loop (GxxGxGK(T/S)) (Figure 6B). Such conserved sequences and structures suggest common evolutionary origins of the ATP-binding P-loop (GxxGxGK(T/S)) and the NAD^+^/NADP^+^- and NADPH-binding GxxxGxG Rossmann domain. These findings are supported by previous reports that P-loop NTPases and Rossmann enzymes with key similar structural elements share a common ancestral origin although they lack apparent homologous sequences^58^. Taken together, our data provide strong structural evidence that N_2_-fixation and photosynthesis catalysts with ATP-binding and NAD^+^/NADP^+^- and NADPH-binding domains share a common protein ancestor which has diverted to present nitrogenase-like proteins, RDH and LPOR during their evolution.

### Phototrophic metabolisms drive the COR to RDH to LPOR evolution

Phototrophic metabolism converts light energy to chemical energy and is one of the main driving forces for evolution in photosynthetic organisms. COR, crucial for bacterial chlorophyll biosynthesis, has played a significant role in this process^59,60^. On the other hand, RDH is a key enzyme in the biosynthesis of retinal that binds to proteins such as rhodopsin in the retina to form photoreceptor proteins, and is associated with vision in animals^61,62^. Previous studies demonstrated that energy transfer from light-harvesting ketocarotenoids to xanthorhodopsins (XRh) occurs in a highly halophilic bacterium and a terrestrial cyanobacterium named *Gloeobacter violaceus*^63,64^. In addition, bacteriorhodopsin (BRh), another vital photoreceptor in halophilic archaea, plays a crucial role in light energy conversion *via* capturing photons to pump protons across the membrane to generate proton motive forces^65^. Both XRh and BRh provide bacteria a simple and efficient energy transfer and conversion system for their adaptation to extreme environments^65,66^. Later, animals have evolved rhodopsins to support vision, while losing photosynthetic functions^67^. Compared to microorganisms, the limited populations and slower reproduction rates of animals have maintained a higher degree of conservation in RDHs. This helps explain why LPOR is most similar to animal rather than prokaryotic RDHs. These led us to question the nature of the evolutionary force that drives the transition from COR to the intermediate protein RDH, and ultimately to LPOR. The evolution of key enzymes in biosynthesis of three energy-converting pigment molecules, chlorophyll-a, bacteriochlorophyll-a, and retinal, directly drives the evolution of photopigments, posing significant implications for life itself. To link all three known phototrophic metabolisms, we propose an evolutionary trajectory for light-dependent metabolism based on our COR®RDH®LPOR evolution analysis (Figure 6C).

The evolution of photosynthetic bacteria was fundamentally driven by their ability to harvest higher redox potentials from light energy, thereby reducing their dependence on specific electron donors. Initially, the anoxygenic photosynthetic bacteria such as Chlorobi, relied on type I reaction center (RCI) and bacteriochlorophyll, using H_2_S as the electron donor (Figure 6C)^68^. Bacteriochlorophyll receives electrons, which are then transferred to RCI to complete photochemical reactions. However, the scarcity of H₂S has led to the evolution of type II reaction centers (RCII), enabling organisms such as Chloroflexota to use organic molecules like acetate as electron donors (Figure 6C)^69^. A newly discovered species within the Chloroflexota phylum utilizes RCI for light energy conversion, despite belonging to a phylum in which RCII is commonly used for phototrophy^69^. Phylogenomic evidence indicates that RCI- and RCII-utilizing *Chloroflexia* share a most recent common phototrophic ancestor, suggesting that RCII has likely evolved within the Chloroflexota phylum (Figure 6C)^69^. Moreover, form I Rubisco in cyanobacteria was thought to originate from Chloroflexota, reinforcing the pivotal role of this phylum in the evolution of photosynthetic mechanisms^70^. Recent findings suggest that cyanobacteria likely evolved from non-photosynthetic bacteria classified as *Sericytochromatia*, into which photosynthetic reaction centers were horizontally transferred^42^. Combining these studies, it is plausible to suggest that *Sericytochromatia* acquired a large set of photosynthesis-related genes, including those for reaction center and form I Rubisco, from Chloroflexota through horizontal gene transfer. This gene acquisition ultimately facilitates the evolution of cyanobacteria ancestor. However, the ancestor of cyanobacteria remains unknown.

Previous studies suggest that primitive Photosystem II (PSII), such as those found in *G. violaceus*, may have originated from a photomanganotroph that utilized Mn²⁺ as an electron donor, leading to the oxidation and formation of high-valent manganese oxides resembling birnessite-type minerals (Figure 6C)^71^. This process likely preceded the full development of the complex mechanisms necessary for efficient oxygen production, and *G. violaceus* may have retained some features of this primitive metabolic pathway (Figure 6C). Early archaean origin of heterodimeric Photosystem I (PSI) in cyanobacteria which split the PsaA and PsaB subunits, occurred over 3.4 billion years ago, preceding the GOE^72^. The integration and evolution of PSII and PSI allowed the ancestor of cyanobacteria to use water as an electron donor and produce oxygen as a byproduct (Figure 6C)^73–75^. As photosynthetic organisms containing oxygenic photosynthesis widely expanded, the oxygen levels on Earth rose rapidly, leading to a decline in the activity of anaerobic enzymes like DPOR and COR and the reduction of bacteriochlorophyll biosynthesis rates. Under the pressure of oxygen-rich environment, COR evolved into RDH, a key enzyme in synthesizing retinal, through genetic recombination thus establishing an oxygen-tolerant phototrophic metabolism system (Figure 6C). Due to the loss of COR, the ancestor of cyanobacteria lost the ability to produce bacteriochlorophyll and had to rely on chlorophyll and retinal as the pigment molecules. Subsequently, RDH acquired an extra region, giving rise to LPOR, an oxygen-tolerant and efficient enzyme (Figure 6A). LPOR marked the emergence of cyanobacteria, reinforcing the dominance of the PSI and PSII systems, which gained a competitive advantage in the evolutionary race. Although RDH was lost, some primitive cyanobacteria, such as *G. violaceus*, retained the ability to carry out phototrophic metabolism using XRh, albeit with carotenoids as the utilized pigments instead of retinal. The photosynthetic activity of cyanobacteria contributed to a significant increase in atmospheric oxygen levels, culminating in the GOE. GOE further consolidates the dominance of phototrophic metabolisms in cyanobacteria, which utilize LPOR to catalyze the biosynthesis of chlorophyll *a* for photosynthesis (Figure 6C). In subsequent evolution, some eukaryotes inherited the photosynthetic system and key chlorophyll-synthesizing enzymes DPOR and LPOR from cyanobacteria, which evolved into chloroplasts (Figure 6C). The chloroplasts ultimately give rise to the most powerful higher autotrophs—plants. Eventually, in flowering plants, DPOR was lost, retaining only the more efficient LPOR. Another group of eukaryotes inherited a retinal-based rhodopsin phototrophic system and key retinal-synthesizing enzyme RDH from the ancient ancestors of cyanobacteria (Figure 6C). Over time, this system gradually lost its function in energy metabolism and may have been repurposed for retinal metabolism in vision systems.

### Design of FeMoco-loaded engineered LPOR protein

Based on the structural insights into nitrogenase-like proteins (DPOR and COR) and SDR-like proteins (RDH and LPOR), we exploit structural cues to design new light-driven electron transfer proteins. In our previous studies, we identified the substrate binding mode in the Pchlide-NADPH-LPOR complex and explored the reaction mechanism using quantum-chemical methods^10,76^. These studies revealed how the LPOR active site can promote the photo-driven reduction through the initial electron transfer from Tyr189 to the substrate, followed by hydride transfer from NADPH and internal electron-transfer from Cys222 to the deprotonated Tyr189 radical^76^. Given the structural and functional similarities between nitrogenase-like proteins and LPOR, particularly the alignment of substrate binding sites, we hypothesize that FeMoco could be accommodated within the LPOR scaffold, enabling light-driven electron transfer to FeMoco, thereby possibly obviating the need for ATP-driven electron transfer to FeMoco observed in all extant natural nitrogenases.

Since the key electron-transfer event that initiates catalysis by LPOR is made possible, in large part, by the active-side amino acids themselves, rather than only the interplay of NADPH and PChlide molecular orbitals, we postulate that these active site features might be sufficient to enable light-induced electron transfer to chemical entities other than Pchlide. Considering that nitrogenase and DPOR are structurally related and that the substrate Pchlide is found to be almost exactly in the position where FeMoco is located in nitrogenase complex^33^, we assume that the LPOR scaffold might be able to accommodate FeMoco and catalyze the light-driven electron-transfer from Tyr189 to the FeMoco.

FeMoco is covalently linked to nitrogenase through a His-Molybdenum bond and a Cys-Fe bond with the cofactor. In LPOR, the substrate binding cavity is partially occluded by a highly mobile loop, which contains a His residue that we decided to co-opt for FeMoco binding. Since no Cys exists at an appropriate position of the bottom of the cavity to bind the opposite end of FeMoco, we changed Thr141 to Cys in our model. By also removing amino acids adjacent to the candidate His in the mobile loop, we thus obtained a modified model of LPOR, designated FeMoco-loaded engineered LPOR (FL-LPOR) hereafter, with His and Cys residues at appropriate distances to bind FeMoco (Figure 7A).

**Figure 7.**
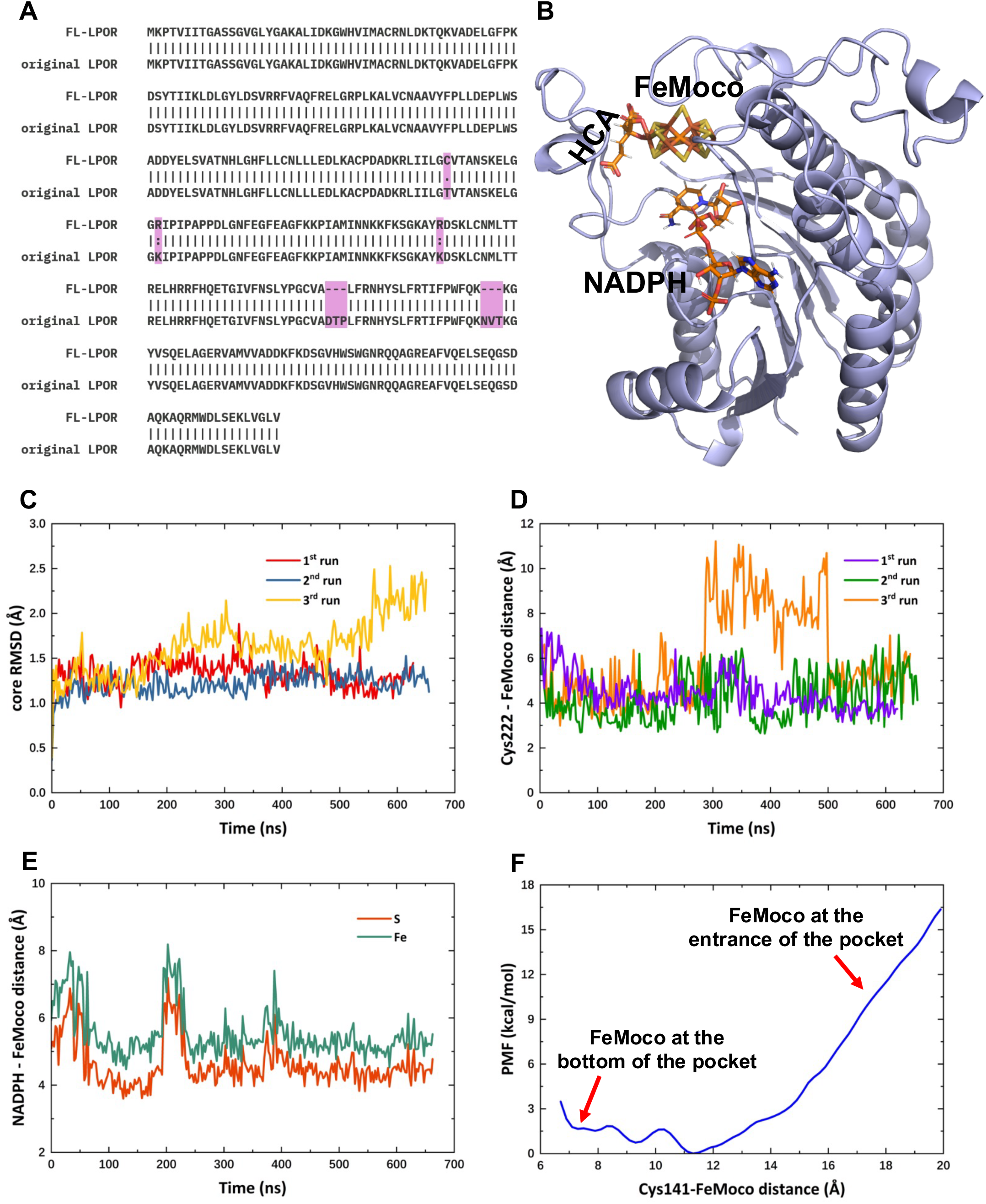
Protein design of FL-LPOR for light-driven nitrogen fixation. (A) Amino acid alignment of naturally occurring LPOR and designed FL-LPOR. (B) A representative structure of FL-LPOR from MD simulation. *R*-homocitrate (HCA) ligand promotes nitrogenase activity by activating the FeMoco, enabling it to reduce dinitrogen under ambient conditions. (C) RMSD of the core portion (i.e. without the flexible lid and C-terminal) of the FL-LPOR along three different molecular simulations. (D) Cys222-FeMoco distance changes along three simulations. (E) Distance between the hydride-donating carbon in NADPH and the S (or Fe) atoms of FeMoco in engineered LPOR along one simulation. (F) Energy profile of cofactor movement from the solution to the active site of engineered LPOR.

Finally, to better mimic the Arg-rich environment provided in naturally occurring nitrogenase to its FeMoco cofactor, two Lys residues were replaced with Arg in FL-LPOR. The modeled structure of FL-LPOR is similar to that of LPOR, allowing for molecular docking with FeMoco at positions comparable to those in LPOR (Figure 7B). Such structure was then submitted to triplicate molecular dynamics simulations (>500 ns each) to test whether FL-LPOR would be sufficiently stable to enable its possible use as a light-driven nitrogen fixation system, in a quest for engineering a possible light-utilizing nitrogenase (LUN) ^34,77^. The results of these simulations show that the bulk of the FL-LPOR remains stable, and that the mobile loop and C-terminal portions retain their high mobility (Figure 7C). The distance between FeMoco and Cys222 (which we predicted to have a crucial role in electron transfer to the substrate) remains short throughout the simulations (Figure 7D), suggesting a reliable pathway for electron transfer. However, the distance between NADPH and FeMoco is more variable. The introduced amino acids appear to have increased the flexibility of the nicotinamide portion of NADPH, which only remained consistently close to FeMoco in one of the three simulations (Figure 7E). These data suggest that while the general binding of FeMoco is stable, the interaction with NADPH might require further optimization to ensure consistent hydride donation to the FeMoco cage. Furthermore, potential of mean force (PMF) graphs depicting FeMoco migration from the entrance of the apo-LPOR cavity to the bottom of the active site suggest that FeMoco entry into the LPOR cavity is thermodynamically favorable (Figure 7F and Video S3). The energy change for FeMoco movement into the cavity is negative, indicating a spontaneous binding process. This suggests that engineered LPOR can readily accommodate FeMoco, which is crucial for enabling subsequent catalytic reactions.

**Video S3.**
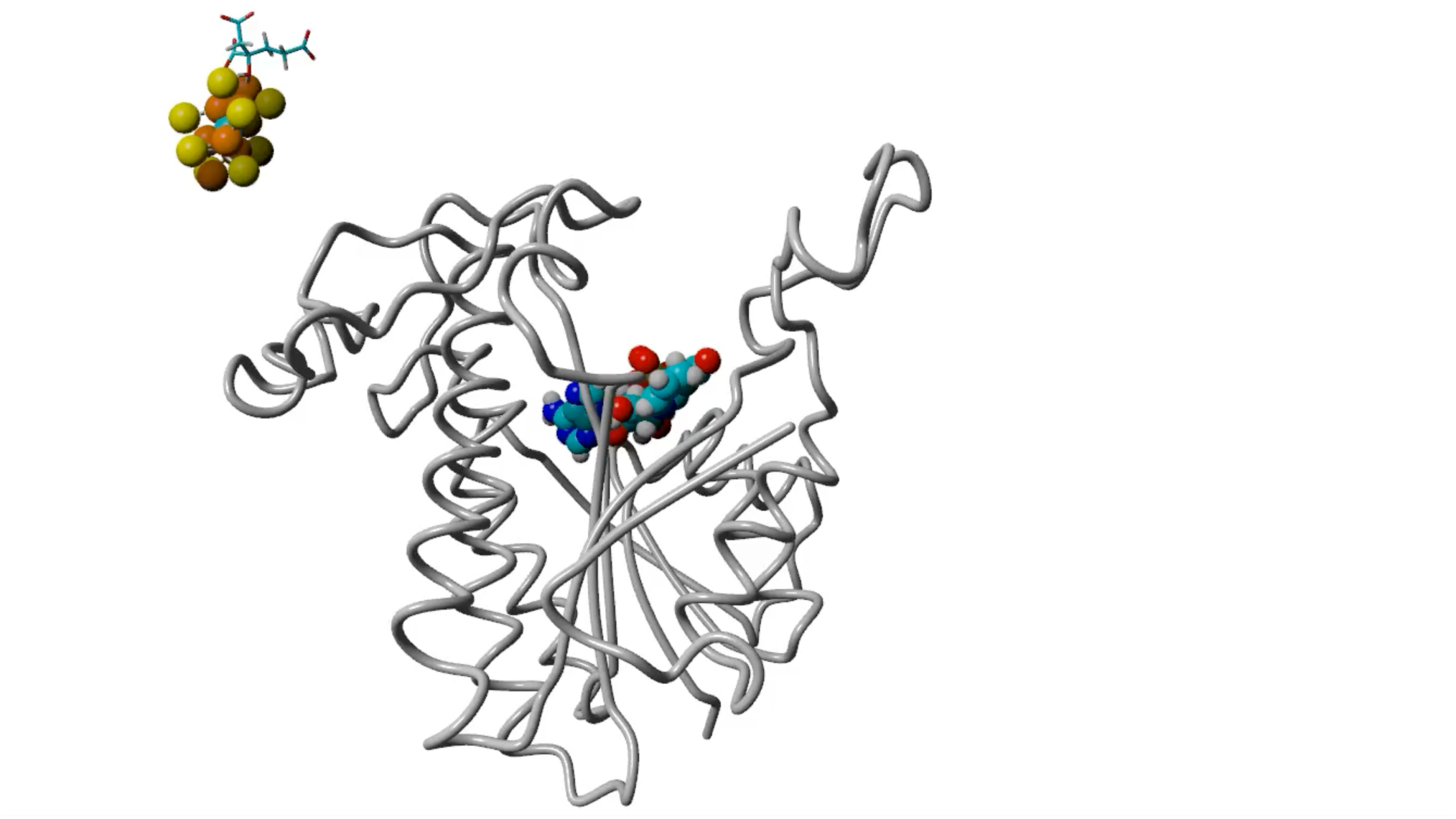
FeMoco integration into the engineered LPOR, related to Figure 7F. The engineered LPOR protein structure was rendered as a pipe model using YASARA to clearly visualize ligand movement. The Fe and S atoms in FeMoco are depicted as orange and yellow spheres, respectively. HCA is depicted as a stick model positioned close to FeMoco. NADPH is depicted as solid space-filling sphere.

## DISCUSSION

Over the past three decades, considerable progress has been achieved in solving crystal structures of nitrogenase, nitrogenase-like protein structures including DPOR, and NADPH-LPOR protein complex. However, the evolutionary relationship among these functionally related proteins remains unexplored. In this study, we uncover how a porphyrin reduction catalyst evolves from a low-efficient ATP-driven DPOR to a highly elevated version of light-driven LPOR. LPOR shares structural similarities with nitrogenase, DPOR and COR subunits, and they all adopt a similar structural fold. Our innovative structure-based analyses indicate that all subunits of nitrogenase-like proteins originate from the CfbC subunit of an ancient nitrogenase-like protein F430. In addition, LPOR evolves from the BchY subunit of multi-subunit COR *via* an unexpected intermediary RDH following genetic recombination and duplication events. This is evidenced by the designed CBRL and CBLL which share structure similarity with RDH and LPOR, respectively. Such COR to RDH to LPOR evolution is consistent with the evolution of phototrophic metabolisms for the synthesis of three energy-converting pigments chlorophyll-a, bacteriochlorophyll-a, and retinal (for vision). Moreover, our findings can be extended to guide the design of a wide range of catalysts, such as a highly efficient light-utilizing version of nitrogenase and photosynthesis with bacteriochlorophyll using light-driven COR (LCOR), leading to the development of more effective forms of BNF and photosynthesis.

### Origin of nitrogenase-like proteins

The query over the origin of nitrogenase and nitrogenase-like proteins associated with the BNF evolution remains intriguing and enigmatic. Through structure-based analyses, we now provide an unambiguous answer to this long-standing question in biology. All nitrogenase-like proteins, including nitrogenase, have evolved from a one-subdomain precursor, i.e. the component 2 subunit CfbC of the ancestral F430 Fe-protein (Figure 5). This subunit underwent duplication(s) and recombination(s) to form multi-subunit complexes crucial for BNF and later for chlorophyll biosynthesis (photosynthesis). The marked similarities in structural folds, particularly the Rossmann fold, across different nitrogenase-like proteins underscore their co-origins and co-functional adaptations. The higher degree of structural similarity among the subunits within component 2 may also account for why nitrogenase retains partial functionality even with the substitution of Fe-protein^38^. Interestingly, we notice that the only two-subdomain subunit CfbD of F340 may represent a unique element in this evolutionary pathway. CfbD seems to play a crucial role in bridging the gap between one-subdomain and three-subdomain precursors during early evolution due to its significant structural similarities and clusters with subunits of other nitrogenase-like proteins (Figure 3). These discoveries further consolidate the notion that all subunits of nitrogenase-like proteins comprise subdomains and share a common evolutionary history. Thus, it is plausible to conclude that the complex nitrogenase-like proteins have evolved from a single simple ancestor through a series of domain duplication and genetic recombination events.

### Origin of LPOR

LPOR is often perceived evolutionarily unrelated to and DPOR. Indeed, LPOR and DPOR lack direct amino acid sequence identity. They belong to different enzyme families. Plant nuclear-encoded LPOR is a light-driven single-chain enzyme, while DPOR, like structurally similar nitrogenase, is an ATP-driven multi-subunit enzymatic complex^31,41^. Physiologically, DPOR is extremely sensitive to oxygen and has existed in early Earth’s anaerobic environments. In contrast, LPOR is oxygen-tolerant and only emerges after atmospheric oxygen level has increased^41^. Nevertheless, a TFT motif was found in DPOR and some LPORs although this motif is absent from the cyanobacterial LPOR, suggesting that the TFT motif in certain LPORs did not originate from DPOR, but could result from convergent evolution^1,72^. On the other hand, LPOR and DPOR both catalyze the reduction of protochlorophyllide to chlorophyllide, despite involving distinct mechanisms under aerobic vs anaerobic conditions in chlorophyll biosynthesis, respectively^9^. Moreover, DPOR can recuse red-light dependent photosynthetic deficiency in chlorotic dormant cyanobacteria^78^. Such similar “LPOR vs DPOR” functionality contradicts the perception of their evolutionary unrelatedness. To resolve this discrepancy, we took an unprecedented structure-based approach to analyze LPOR evolution. Instead of pairwise-comparing protein sequences, we conducted an in-depth regional and global structural similarity study and discovered extensive continuous structural similarities between LPOR and nitrogenase-like proteins including DPOR and COR far exceeding the length of the TFT motif. The overall regions with structural similarity cover over 71% of entire LPOR (Table 1). Surprisingly, LPOR was found to be more closely related to COR than DPOR. Such similar structures may result from recombination-driven events, as inferred in previous structural comparisons^55^. Thus, the “COR/DPOR to LPOR” is less likely to be coincidental, but more probably rooted in evolutionary reality.

Our structural analysis identified critical β-α-β segments that underpin the evolution of nitrogenase-like proteins, RDH, and LPOR, showcasing a conserved structural continuity from component 2 to LPOR (Figure 6A and 6B). This supports previous hypothesis that both P-loop NTPase and Rossmann folds share a β-α-β ancestral fragment^58^ and extends it by revealing the structural elaboration of this fragment during COR-to-LPOR evolution. By tracing the origins of domains involved in ATP-, NAD^+^/NADP^+^-, and NADPH-dependent catalysis, our study provides structural and functional evidence for the shared ancestry of nitrogenase-like proteins, RDH, and LPOR. It further highlights the evolutionary adaptation of these folds for distinct biochemical roles, shedding new light on the origins of nitrogen fixation and photosynthesis catalysts.

Moreover, we uncovered significant sequence and structural similarities between LPOR and proteins in the SDR family. This suggests that LPOR shares a common ancestor with SDR proteins, of which the closest match to LPOR is RDH. Phylogenetic analyses also link LPOR with other SDR proteins^4^. Collectively, our structure-based analyses unravel a previously unknown “COR/DPOR to RDH to LPOR” evolutionary path. LPOR likely evolves from an SDR family member, i.e. RDH as an essential intermediary protein, through domain shuffling and genetic recombination. Our work also demonstrates that comparative structural analysis can be an important tool in linking sequence-unrelated proteins and alike functionality, revising and/or re-defining well-established evolutionary theories.

### Evolutionary relatedness between BNF and photosynthesis

BNF and photosynthesis associated with nitrogen- and carbon-cycle are two of the most important processes for life to survive and thrive on Earth. BNF, nitrogenase and nitrogenase-like proteins likely evolved in early Earth history when the biosphere lacked oxygen before the rise of LPOR-mediated oxygenic photosynthesis in cyanobacteria, and later in land plants^77,79^. Previous studies, together with our work described here, clearly demonstrate that nitrogenase and nitrogenase-like proteins including DPOR and COR are of a common origin.

Furthermore, LPOR is now found to have evolved from a common ancestor shared with nitrogenase and other related reductases. Therefore, the co-origin of these key enzymes in photosynthesis and BNF reveals for the first time the clear co-evolutionary relationship between the two physiological processes, which can still be simultaneously impacted by genetic and epigenetic mutations in modern crops^80^.

### Evolutionary link between plant photosynthesis and animal vision

Our findings suggest a significant evolutionary link between plant photosynthesis and animal vision (Figure 6C). RDH in animals shows significant sequence and structural similarities with LPOR. This indicates a potential evolutionary transition from plant phototrophism (photosynthesis) to animal visual systems. RDHs are crucial for synthesizing retinal, which combines with opsins to form rhodopsin, the photoreceptor protein in animal retina^61,62^. Many modern eye types emerged rapidly during the Cambrian period, about 530 million years ago, coinciding with appearance of land plants approximately 450 million years ago^81,82,79^. The high sequence and structural conservation between animal RDHs and plant LPORs suggests these enzymes have retained their functional roles over billions of years. Such conservation highlights possible retinal-based photoreception in photosynthesis and vision.

The possible evolutionary transition from COR to LPOR is facilitated by RDH, and the latter might have set the stage for developing sophisticated animal visual systems. This unexpected connection underscores a deep evolutionary link between light absorption mechanisms in plants and vision development in animals. Eukaryotic organisms may have engulfed different cyanobacteria and utilized them for the benefit of development and evolution. For example, chloroplasts develop after plant engulfing cyanobacteria, retaining DPOR and LPOR-driven photosystems. It is possible that vision cells develop and evolve key components like retinal and rhodopsin by a similar approach in animals^83,84^.

### The key role of phototrophic metabolism in evolution

Our analysis suggests that the formation of LPOR underwent two key recombination events (Figure 5, Video S2). First, the recombination of BchX and BchY generated an intermediate form of LPOR. Subsequently, an extra region from an unknown source was incorporated into this intermediate protein, forming the primitive LPOR. Structural similarity analyses between LPOR and COR indicate that the LPOR intermediate likely contained RDH, which provided a critical evolutionary link between COR and LPOR. The increase in oxygen concentration resulting from the water oxidation activity of PSII exerted significant selective pressure, shaping the evolution of phototrophic metabolism. Evidence from primitive cyanobacteria like *G. violaceus* supports a transitional phase in energy metabolism between photosynthetic bacteria and cyanobacteria using retinal as a photosynthetic pigment^63,64^. The role of RDH in retinal-mediated phototrophic metabolism and its eventual transition to LPOR highlights the adaptability of biological systems to new environmental pressures and energy demands. The evolution of LPOR from COR *via* RDH reflects a strategy of utilizing existing genetic materials and reshuffling domains to efficiently develop new functionalities, highlighting the significant role of phototrophic metabolism in driving evolutionary changes and enhancing the efficiency of biological systems.

### Life evolution single-plank bridge

The emergence of photosynthesis profoundly reshaped the evolutionary trajectory of life on Earth, providing energy sources for ecosystems and significantly elevating atmospheric oxygen levels, thereby laying the foundation for the proliferation of aerobic organisms^85,86^. During this process, a series of key enzymes, ranging from nitrogenase-like proteins (DPOR/COR) to SDR proteins (LPOR), played pivotal roles (Figure 6A and 6C). Among them, LPOR is crucial in the rate-limiting step of chlorophyll biosynthesis, enabling proton and hydride transfer on the sub-nanosecond timescale and thus providing the molecular basis for efficient chlorophyll formation^87,88^. By drastically enhancing the ability of life on Earth to harness solar energy, LPOR overcame the energy shackles in species evolution, which arose from the vast energy demands imposed by the complexity and sheer size of macroscopic organisms.

Based on our analysis of the COR→RDH→LPOR transition, we propose the key enzymes-based theory of the “single-plank bridge” in life evolution, which represents a crucial pathway that enabled primitive life (microorganisms) to overcome the energy shackles and paved the way for the emergence of higher organisms (macroscopic organisms) (Figure 8). In this evolutionary framework, these enzymes can be traced back to the last universal common ancestor (LUCA), from which the major domains of bacteria and archaea diverged^89,90^. The early bacterial and archaeal lineages evolved distinct metabolic traits, including F430 and nitrogenase, establishing foundational bioenergetic pathways. Over time, bacterial branches evolved COR and DPOR, which conferred rudimentary photosynthetic capabilities. LPOR and RDH originated from COR through gene rearrangement and horizontal transfer, gaining novel functions that supported phototrophic metabolism (Figure 6A). These key enzymes—particularly RDH and LPOR/DPOR—into primitive eukaryotic lineages (Fungi/Protozoa) via symbiosis or horizontal gene transfer provided critical advantages in energy acquisition (photosynthesis) and environmental sensing (vision) (Figure 6C and 8). Thus, these enzymes, spanning from microbial predecessors to macroscopic plants and animals, effectively link the earliest phases of life’s origin with the complex biosphere we recognize today (Figure 8). By evolving key enzymes like LPOR and RDH, this “single-plank bridge” connects photosynthesis in plants and vision in animals, underscoring the deep interdependence between autotrophs and heterotrophs in Earth’s ecosystems. Plants supply oxygen and food, while animals aid in seed and pollen dispersal (Figure 8). These interactions highlight how phototrophic metabolism and its innovations (e.g., LPOR and RDH) have collectively shaped the co-evolution of life on Earth (Figure 8). This theory offers a unified perspective on the evolutionary pathways that led to the rapid diversification of higher organisms and also provides valuable insights for astrobiology in the search for extraterrestrial macroscopic life forms.

**Figure 8.**
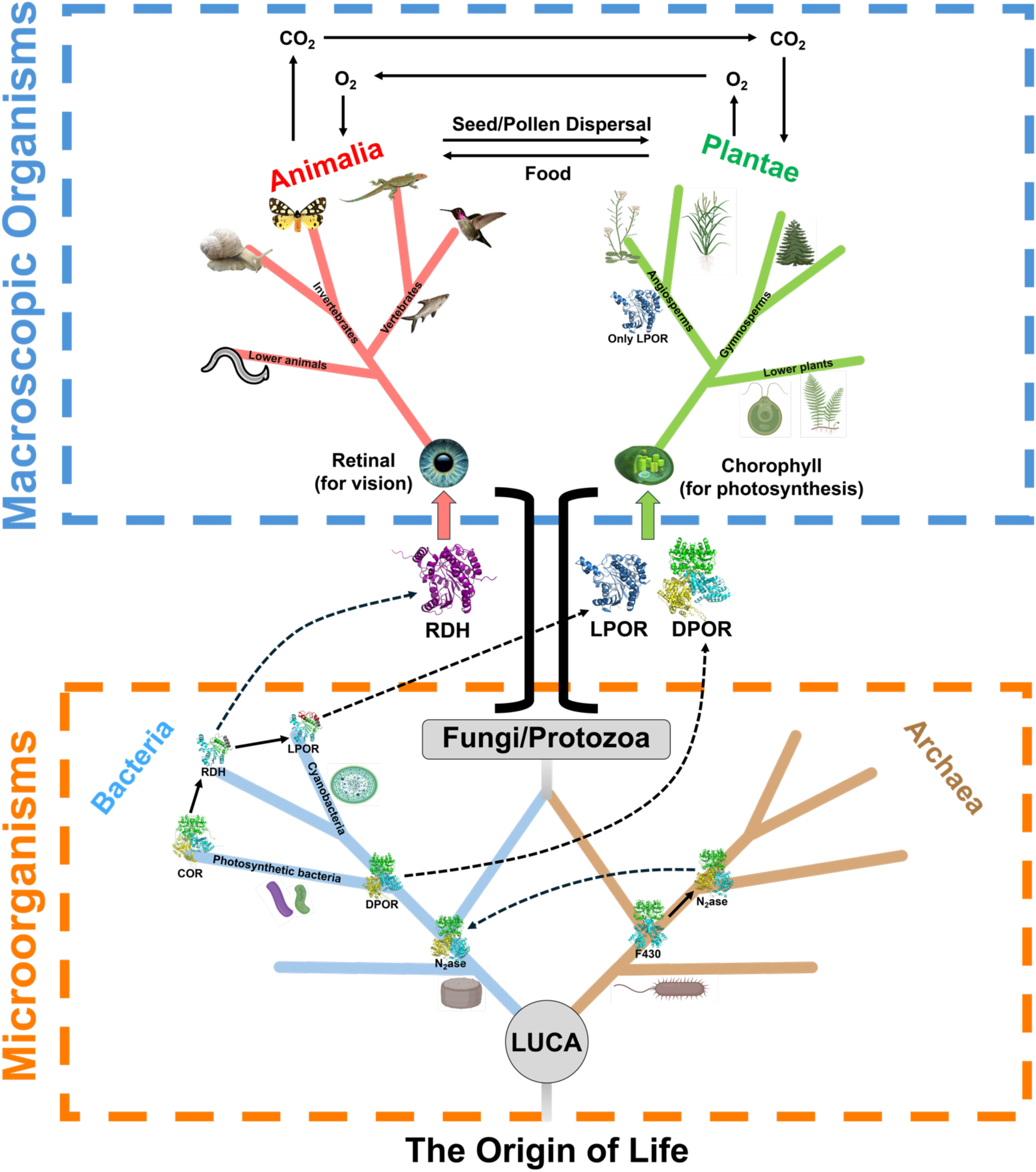
The “single-plank bridge” in life evolution. This key enzymes-based theory posits that the evolutionary path of nitrogenase-like proteins and SDR proteins, particularly the COR®RDH®LPOR transition, represents a pivotal evolutionary juncture. The emergence of LPOR enabled the efficient utilization of solar energy via photosynthesis, providing the energetic foundation for the evolution of complex life. Concurrently, RDH played a crucial role in retinal biosynthesis, driving the development of Animalia’s vision system. LPOR and RDH bridged the evolutionary transition from microorganisms (LUCA, bacteria and archaea) to macroscopic organisms (Plantae and Animalia), overcoming the energy shackles barrier and forging a link to vision.

### Practical applications and prospects

The insights gained from this research not only deepen our understanding of key enzymes associated with BNF and photosynthesis, but also have significant implications for designing novel biocatalysts. One promising application is developing a LUN,bypassing many limitations of nitrogenase’s oxygen sensitivity and ATP dependence^34,77^. By leveraging insights from LPOR and nitrogenase, we can engineer a novel nitrogenase that harnesses light energy to fix nitrogen, potentially increasing BNF efficiency (Figure 9A). Unlike traditional nitrogenase systems confined to leguminous plants with root nodules (Figure 9A), LUN systems aim to decouple nitrogen fixation from symbiotic relationships, offering a scalable solution for non-leguminous crops. For instance, incorporating LUN into crops like rice, wheat, and maize, which currently depend heavily on chemical fertilizers, could reduce fertilizer reliance by up to 50% while increasing crop yields in nitrogen-limited soils. Such innovations align with global goals for sustainable agriculture, particularly in regions with high nitrogen demand. Implementing this in oxygen-rich and eukaryotic environments presents additional hurdles due to FeMoco’s oxygen sensitivity. Ongoing efforts are focused on designing oxygen-tolerant FeMoco variants^91–93^.

**Figure 9.**
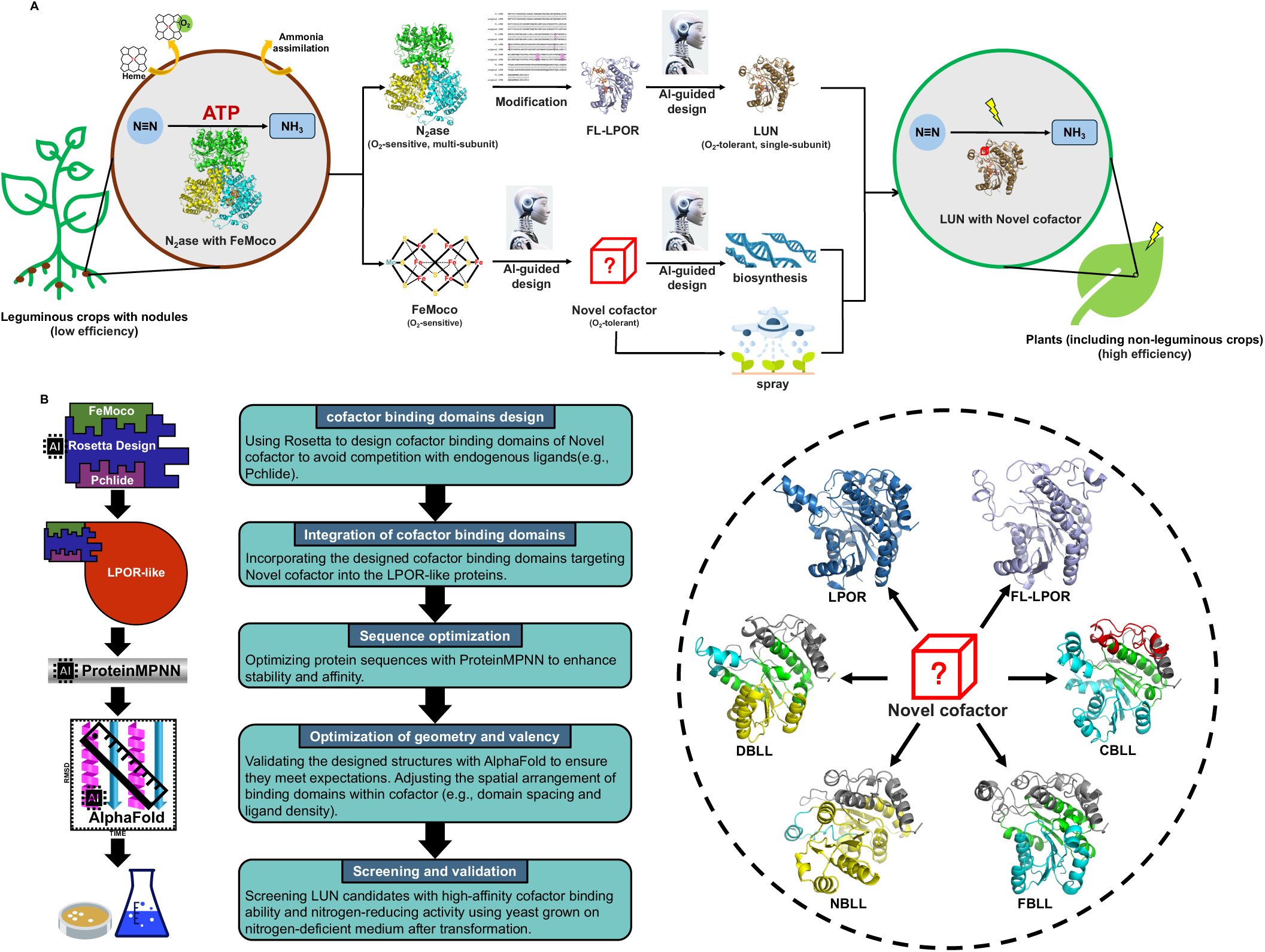
Engineering roadmap and AI-guided design pathways for LUN in future. **(A)** Engineering roadmap a LUN system with enhanced N_2_-fixation efficiency. Traditional anaerobic FeMoco-containing nitrogenase, which consumes ATP, is typically found in leguminous plants and exhibits low efficiency. The proposed modification utilizes AI-based design to develop a modified FeMoco and a LUN system. It involves modifications of FL-LPOR and the design of a novel cofactor that functions aerobically, potentially increasing efficiency of nitrogen fixation in all plants. The alignment shows the modification to LPOR to enable it to bind FeMoco. **(B)** AI-based design of LUN with enhanced N_2_-fixation efficiency. The process begins with cofactor binding domain design using Rosetta to target novel cofactors, avoiding competition with endogenous ligands. The designed domains are then integrated into LPOR-like proteins. Sequence optimization is performed with ProteinMPNN to enhance stability and affinity. The geometry and valency of the designed structures are initially predicted using AlphaFold, followed by molecular dynamics simulations for validation and optimization of spatial arrangements and ligand density. Finally, the screening and validation step involves selecting LUN candidates with high-affinity cofactor binding ability and nitrogen-reducing activity through yeast cultivation on nitrogen-deficient medium after transformation.

Based on the successful computational protein design strategies developed by David Baker’s group^94,95^, the development of an AI-guided LUN begins with the design of cofactor binding domain using Rosetta to optimize compatibility with novel cofactors, minimizing interference with endogenous ligands such as Pchlide (Figure 9B). Following this, integration of the designed cofactor binding domains into LPOR-like proteins establishes candidates for efficient light-driven catalysis. Furthermore, AI-based sequence optimization with ProteinMPNN ensures enhanced protein stability and affinity. Validation of the structural arrangement of the cofactor binding domains, including spatial configuration and ligand density, can be initiated with AlphaFold for structure prediction, followed by molecular dynamics simulations to ensure precise geometric and valency adjustments (Figure 9B). Finally, screening and validation steps involve testing LUN candidates with high-affinity cofactor binding ability and nitrogen-reducing activity using yeast grown on nitrogen-deficient media after transformation (Figure 9B).

This innovation could revolutionize agriculture by reducing reliance on chemical fertilizers and enhancing sustainable farming practices. Moreover, the further analysis of sub-domains and the exploration of effector proteins have provided a methodology for the engineering design of enzyme proteins. Most importantly, this study offers new theoretical frameworks for future research on BNF engineering. Similarly, the concept of a LCOR opens new avenues for enhancing photosynthetic efficiency. Designing new LPOR-like enzymes could enhance chlorophyll synthesis and photosynthetic efficiency for enhancing crop productivity.

### Limitations of the study

#### AI-assisted design of LUN remains a challenge

This study presents a theoretical framework suggesting the evolutionary transition from multi-subunit nitrogenase-like proteins to single-subunit LPOR-like proteins, proposing that this transition could be experimentally validated through the artificial design of a LUN. The successful design of such a LUN would serve as strong evidence supporting our evolutionary theory. Significant challenges may hinder this validation process: FeMoco might not fit the active site on engineered LPOR, its binding might not be spontaneous, and the FL-LPOR might not be structurally stable. Additionally, FeMoco is oxygen-sensitive, which will render the FL-LPOR construct liable to degradation by oxygen and prevent its straightforward utilization in oxygen-evolving photosynthetic environments. The current limitations in AI-driven enzyme design, particularly in creating complex enzymes like LUN, make this task difficult. Additionally, the experimental validation of LUN would require the synthesis or modification of FeMoco, a complex cofactor essential for nitrogenase function. With the development of AI technologies, particularly in designing complex enzymes and cofactors, we anticipate that this evolutionary theory will be thoroughly tested and validated.

Our molecular dynamics simulations provide promising insights into the stability of FeMoco within the engineered LPOR, and show that the overall stability of the protein and the favorable energy profile for FeMoco binding indicate that this system has potential as a light-driven nitrogen fixation catalyst. Further refinement may be needed to stabilize the NADPH-FeMoco interaction, thereby enhancing the efficiency of the putative hydride transfer mechanism from NADPH to the cofactor during N_2_ catalysis. Two major challenges remain, the system’s sensitivity to oxygen and the lack of understanding regarding the feasibility of proposed light-driven electron transfers from the amino acids of engineered LPOR to FeMoco. The first challenge poses a significant barrier to practical applications, but may be circumvented by expressing the *FL-LPOR* in organisms like *Anabaena*, since these photosynthetic organisms are able to differentiate into non-oxygen evolving heterocysts whenever they need to generate an anaerobic intracellular environment for nitrogen fixation. The second challenge does not seem, at present, amenable to computational simulation due to the overwhelming complexity of the electronic and chemical character of FeMoco (with dozens of possible electronic and protonation states), and must therefore await experimental verification through the expression of *FL-LPOR* in FeMoco-synthesizing organisms.

## STAR★METHODS

### Structural modeling

LPOR structures and non-natural proteins were predicted using AlphaFold v2.3.1^96^ in monomer mode with the full_dbs database, which was downloaded using a script from DeepMind on July 20, 2023. For each protein, five models were generated (model_1, model_3, model_4, model_5, and model_2_ptm) to obtain the pTM score. The best model (ranked_0.pdb) was selected based on the average pLDDT score. The structures of nitrogenase-like proteins were predicted using AlphaFold-Multimer v3, which generated 25 possible protein complex structures using five different models. The accuracy and reliability of these structures were assessed using the ipTM+pTM scoring mechanism, with the best structures (ranked_0.pdb) selected for subsequent analysis^97^.

For proteins with existing crystal structures, unmodeled regions are often present due to insufficient resolution in certain parts of the crystal structure data or structural disorder of some regions during the experiment, making these regions challenging to observe. When a protein had multiple crystal structures, PyMOL (The PyMOL Molecular Graphics System, Version 2.0 Schrödinger, LLC.) was used to align these structures, allowing for the identification of regions unmodeled in one structure but resolved in another. This comparative approach helped integrate information from multiple crystal structures to achieve the most complete protein model. For proteins that were still incomplete after this process, AlphaFold2 was utilized to fill in the gaps, ultimately ensuring a comprehensive structure for subsequent analysis. This methodology is a key component of the approach detailed in this paper. Topological graphs were generated using Pro-origami^98^.

### Structural alignment

The PyMOL software was utilized to dissect the 3D structures of proteins, extracting the structural components of each subunit. Subsequently, the (split) structures were pairwise aligned using the DALI server^99^, generating DALI Z-scores to assess all-against-all structural similarities, and a DALI Z-score with a cutoff >2 indicating structure similarity. To ensure consistency, given that the DALI server may yield varying values for identical alignments across different datasets, the DALI Z-score matrices used in all heatmaps in this study were extracted from a single comparison matrix that involved all proteins. This approach was taken to maintain uniformity across data presentations. The structural similarity relationships were visualized through the structural similarity dendrogram of the DALI server and the heatmap of TBtools^100^. Details of the similar regions were determined using US-align, with similar regions identified as aligned residues^101^. Additionally, PyMOL was employed to analyze the 3D structures of proteins to identify helices and sheets. Based on the information from the structural similarity alignments, structures most homologous to LPOR were identified. The corresponding sequences were then used to generate synthetic protein sequences. Based on structural similarity comparisons and secondary structure regions, the structures exhibiting most homologous to LPOR were identified and combined to generate non-native amino acid sequences.

### Selection criteria for deducing LPOR origin

A distance-based algorithm using US-align was employed to compare structural regions, enabling the identification of structural regions relevant to the evolutionary origin of LPOR. These regions were selected based on two criteria: (1) a distance within 5.0 Å of the target structure, and (2) the closest secondary structure alignment to that of LPOR. When multiple regions met both criteria equally, the longest continuous similar region was chosen. To maintain unique comparisons, regions associated with nitrogenase-like proteins were excluded from subsequent comparisons when feasible. This approach prioritized regions with the highest homology, focusing on shared evolutionary origins rather than merely structural similarity.

### Evaluation systems for ancestral-like LPORs

The evaluation system for ancestral-like LPORs was based on criteria including structural continuity, similarity score, amino acid substitution ratio, and evidence supporting the presence of extra regions. Regions with higher structural continuity, substitution ratio and similarity scores were prioritized. In contrast, additional amino acids in extra regions that did not match LPOR were considered less indicative of shared ancestry. Notably, the similarity scores were influenced by the substitution ratio, with a higher number of substitutions leading to lower similarity scores due to the reduced presence of the original LPOR sequence. When similarity scores were close, sequences with a higher substitution ratio were preferred, as they retained more structural features relevant to LPOR. This method ensures that the most structurally relevant and evolutionarily informative regions are prioritized.

### Molecular docking

Molecular docking studies were conducted using the protein structure of COR generated by AlphaFold2. Full hydrogen atoms were added to the COR structure using MGLTools 1.5.7 (http://ccsb.scripps.edu/mgltools/), and the protein was subsequently prepared as a receptor (Macromolecule). Chlide was selected as the ligand for this study. Using MGLTools 1.5.7, full hydrogen atoms were added to the ligand. Subsequently, charges were automatically assigned to the ligand atoms. Torsion bonds were then detected and those relevant to the ligand’s conformational flexibility were selected within the torsion tree for further analysis. Autogrid4 was employed to define the docking grid box, focusing on the kinase domain. The grid box parameters generated by Autogrid4 were used to create the configuration file for AutoDock Vina^102^ with the assistance of MGLTools 1.5.7. Subsequently, AutoDock Vina was executed to perform the docking simulations, generating the resulting docking data. The FeMoco docked into engineered LPOR was also generated by AutoDock Vina. The molecular docking of NAD^+/^NADP^+^ cofactor into RDH were predicted by AlphaFold3^103^.

### Molecular dynamics simulations

Molecular dynamics simulations were conducted using the AMBER03 force field in YASARA to analyze the stability of FL-LPOR^104,105^. Initial FeMoco-binding position was computed with AutoDock Vina^102^. FeMoco structure was optimized at the PBE0-D3/6-31+G(2d,p) theory level using ORCA 5.0.3^106^. Charges were computed from the wavefunction using the RESP method^107^, as implemented in Multiwfn^108^. The protein was simulated in a cubic cell with dimensions 86 Å along each axis, containing ∼63,800 atoms, with a NaCl concentration of 0.9%. The simulations employed a dual-time strategy: 2.5 fs for intramolecular forces and 5 fs for intermolecular forces. An 8 Å cutoff was applied for Lennard-Jones interactions and electrostatic interactions were calculated using the particle mesh Ewald method^109^. Simulated annealing minimizations started at 298 K, with velocities scaled down by 0.9 every 10 steps for 5 ps. Following annealing, simulations continued at 298 K for at least 500 ns, maintaining temperature control *via* a Berendsen thermostat. Structural stability was assessed by calculating the RMSD (Root Mean Square Deviation) for three regions: the protein core (a.a. 1-225), the flexible C-terminal tail (a.a. 249-280), and the full protein throughout the simulation duration.

### PMF calculations

The potential of mean force (PMF) calculations were performed to evaluate movement into the active site of the mutated LPOR. The PMF was determined using umbrella sampling, where the distance between key residues and FeMoco was constrained using a harmonic potential of the form *V* = 1/2*k*(*x* − *x*_0_)^2^ with a force constant of 8.0 kcal/mol/Å^2^. Sampling was carried out in 0.4 Å intervals, with each interval (or bin) sampled for 6.2 ns. The first 1.2 ns of data from each bin were discarded to ensure proper equilibration. The final PMF profile was obtained using the weighted histogram analysis method (WHAM) ^110^.

## SUPPLEMENTAL INFORMATION

Supplemental information can be found online.

## ACKNOWLEDGMENTS

We thank Alison Smith, Andy Sayer, James W Murray, Xiaoquan Qi, Long Mao, Aiwu Zhou, and Zengtao Zhong for their constructive suggestions. This research was supported by the funds of Ministry of Science and Technology of China (2019YFA0904700) and the National Natural Science Foundation of China (32471477) to C.Q., by the funds of the National Natural Science Foundation of China (32330096) and Natural Science Foundation of Hebei Province (C2024204246) to J.J.Z, and by the “Hundred Talents Program” for the introduction of high-level overseas talents in Hebei Province (E2020100004) and the Chunhui Talent Project Provincial Natural Science Foundation of Hebei Province (C2022204116) to L.M..

## AUTHOR CONTRIBUTIONS

C.Q. conceived the study. X.S., J.J.Z., L.M. and C.Q. designed the analysis and experiment. X.S. and C.M. collected and processed data. X.S., L.M., P.S., and Y.H. analyzed the data. X.S., J.J.Z., L.M., P.S., and C.Q. discussed and interpreted the results. X.S., L.M., and C.Q. propose the theory. X.S., L.M. and C.Q. drafted the original manuscript, and Y.H. helped re-draft the original manuscript with inputs from F.Z., R.D., Z.R., G.Z., J.Z., C.Z., M.W., X.Y., J.Y., S.P., E.E, and N.S..

## REFERENCES

1. Olejarz, J., Iwasa, Y., Knoll, A.H., and Nowak, M.A. (2021). The Great Oxygenation Event as a consequence of ecological dynamics modulated by planetary change. Nature Communications 12. ARTN 3985 10.1038/s41467-021-23286-7.

2. Schirrmeister, B.E., de Vos, J.M., Antonelli, A., and Bagheri, H.C. (2013). Evolution of multicellularity coincided with increased diversification of cyanobacteria and the Great Oxidation Event. Proc Natl Acad Sci U S A 110, 1791–1796. 10.1073/pnas.1209927110.

3. Yamazaki, S., Nomata, J., and Fujita, Y. (2006). Differential operation of dual protochlorophyllide reductases for chlorophyll biosynthesis in response to environmental oxygen levels in the cyanobacterium Leptolyngbya boryana. Plant Physiol. 142, 911–922. 10.1104/pp.106.086090.

4. Yang, J., and Cheng, Q. (2004). Origin and evolution of the light-dependent protochlorophyllide oxidoreductase (LPOR) genes. Plant Biol. 6, 537–544. 10.1055/s-2004-821270.

5. Schirrmeister, B.E., Gugger, M., and Donoghue, P.C. (2015). Cyanobacteria and the Great Oxidation Event: evidence from genes and fossils. Palaeontology 58, 769–785. 10.1111/pala.12178.

6. Nascimento, S.M., Zou, Y., and Cheng, Q. (2016). Review of studies on the last enzymes in bacteriochlorophyll (Bchl) and chlorophyll (Chl) biosynthesis. American Journal of Plant Sciences 7, 1639–1651.

7. Wietrzynski, W., and Engel, B.D. (2021). Chlorophyll biogenesis sees the light. Nat Plants 7, 380–381. 10.1038/s41477-021-00900-6.

8. Heyes, D.J., Zhang, S.W., Taylor, A., Johannissen, L.O., Hardman, S.J.O., Hay, S., and Scrutton, N.S. (2021). Photocatalysis as the ‘master switch’ of photomorphogenesis in early plant development. Nature Plants 7, 268–276. 10.1038/s41477-021-00866-5.

9. Nomata, J., Kondo, T., Mizoguchi, T., Tamiaki, H., Itoh, S., and Fujita, Y. (2014). Dark-operative protochlorophyllide oxidoreductase generates substrate radicals by an iron-sulphur cluster in bacteriochlorophyll biosynthesis. Sci Rep 4, 5455. 10.1038/srep05455.

10. Zhang, S.W., Heyes, D.J., Feng, L.L., Sun, W.L., Johannissen, L.O., Liu, H.T., Levy, C.W., Li, X.M., Yang, J., Yu, X.L., et al. (2019). Structural basis for enzymatic photocatalysis in chlorophyll biosynthesis. Nature 574, 722–725. 10.1038/s41586-019-1685-2.

11. Nguyen, H.C., Melo, A.A., Kruk, J., Frost, A., and Gabruk, M. (2021). Photocatalytic LPOR forms helical lattices that shape membranes for chlorophyll synthesis. Nature Plants 7, 437–444. 10.1038/s41477-021-00885-2.

12. Liu, M., Ma, W., Su, X., Zhang, X., Lu, Y., Zhang, S., Yan, J., Feng, D., Ma, L., and Taylor, A. (2022). Mutation in a chlorophyll-binding motif of Brassica ferrochelatase enhances both heme and chlorophyll biosynthesis. Cell reports 41.

13. Curatti, L., and Rubio, L.M. (2017). Corrigendum to “Challenges to develop nitrogen-fixing cereals by direct nif-gene transfer” [Plant Sci. 225 (August) (2014) 130-137]. Plant Sci. 260, 70. 10.1016/j.plantsci.2017.04.002.

14. Rubio, L.M., and Ludden, P.W. (2008). Biosynthesis of the iron-molybdenum cofactor of nitrogenase. Annu. Rev. Microbiol. 62, 93–111. 10.1146/annurev.micro.62.081307.162737.

15. Garcia, A.K., Kolaczkowski, B., and Kacar, B. (2022). Reconstruction of Nitrogenase Predecessors Suggests Origin from Maturase-Like Proteins. Genome Biol Evol 14. 10.1093/gbe/evac031.

16. Rucker, H.R., and Kacar, B. (2023). Enigmatic evolution of microbial nitrogen fixation: insights from Earth’s past. Trends Microbiol. 10.1016/j.tim.2023.03.011.

17. Jasniewski, A.J., Lee, C.C., Ribbe, M.W., and Hu, Y. (2020). Reactivity, Mechanism, and Assembly of the Alternative Nitrogenases. Chem Rev 120, 5107–5157. 10.1021/acs.chemrev.9b00704.

18. Georgiadis, M.M., Komiya, H., Chakrabarti, P., Woo, D., Kornuc, J.J., and Rees, D.C. (1992). Crystallographic structure of the nitrogenase iron protein from Azotobacter vinelandii. Science 257, 1653–1659. 10.1126/science.1529353.

19. Kim, J., and Rees, D.C. (1992). Structural models for the metal centers in the nitrogenase molybdenum-iron protein. Science 257, 1677–1682. 10.1126/science.1529354.

20. Kim, J., and Rees, D.C. (1992). Crystallographic structure and functional implications of the nitrogenase molybdenum-iron protein from azotobacter vinelandii. Nature 360, 553–560. 10.1038/360553a0.

21. Peters, J.W., and Szilagyi, R.K. (2006). Exploring new frontiers of nitrogenase structure and mechanism. Curr. Opin. Chem. Biol. 10, 101–108. 10.1016/j.cbpa.2006.02.019.

22. Seefeldt, L.C., Hoffman, B.M., Peters, J.W., Raugei, S., Beratan, D.N., Antony, E., and Dean, D.R. (2018). Energy Transduction in Nitrogenase. Acc Chem Res 51, 2179–2186. 10.1021/acs.accounts.8b00112.

23. Howard, J.B., and Rees, D.C. (1996). Structural Basis of Biological Nitrogen Fixation. Chem Rev 96, 2965–2982. 10.1021/cr9500545.

24. Seefeldt, L.C., Hoffman, B.M., and Dean, D.R. (2009). Mechanism of Mo-dependent nitrogenase. Annu. Rev. Biochem. 78, 701–722. 10.1146/annurev.biochem.78.070907.103812.

25. Schindelin, N., Kisker, C., Sehlessman, J.L., Howard, J.B., and Rees, D.C. (1997). Structure of ADP center dot AIF(4)(-)-stabilized nitrogenase complex and its implications for signal transduction. Nature 387, 370–376. DOI 10.1038/387370a0.

26. Howard, J.B., and Rees, D.C. (2006). How many metals does it take to fix N2? A mechanistic overview of biological nitrogen fixation. Proc Natl Acad Sci U S A 103, 17088–17093. 10.1073/pnas.0603978103.

27. Rutledge, H.L., Cook, B.D., Nguyen, H.P.M., Herzik, M.A., Jr., and Tezcan, F.A. (2022). Structures of the nitrogenase complex prepared under catalytic turnover conditions. Science 377, 865–869. 10.1126/science.abq7641.

28. Zheng, K.Y., Ngo, P.D., Owens, V.L., Yang, X.P., and Mansoorabadi, S.O. (2016). The biosynthetic pathway of coenzyme F430 in methanogenic and methanotrophic archaea. Science 354, 339–342. 10.1126/science.aag2947.

29. Sorokin, D.Y., Makarova, K.S., Abbas, B., Ferrer, M., Golyshin, P.N., Galinski, E.A., Ciordia, S., Mena, M.C., Merkel, A.Y., Wolf, Y.I., et al. (2017). Discovery of extremely halophilic, methyl-reducing euryarchaea provides insights into the evolutionary origin of methanogenesis. Nature Microbiology 2. ARTN 17081 10.1038/nmicrobiol.2017.81.

30. Moore, S.J., Sowa, S.T., Schuchardt, C., Deery, E., Lawrence, A.D., Ramos, J.V., Billig, S., Birkemeyer, C., Chivers, P.T., Howard, M.J., et al. (2017). Elucidation of the biosynthesis of the methane catalyst coenzyme F(430). Nature 543, 78–82. 10.1038/nature21427.

31. Vedalankar, P., and Tripathy, B.C. (2019). Evolution of light-independent protochlorophyllide oxidoreductase. Protoplasma 256, 293–312. 10.1007/s00709-018-1317-y.

32. Ghebreamlak, S.M., and Mansoorabadi, S.O. (2020). Divergent Members of the Nitrogenase Superfamily: Tetrapyrrole Biosynthesis and Beyond. ChemBioChem 21, 1723–1728. 10.1002/cbic.201900782.

33. Muraki, N., Nomata, J., Ebata, K., Mizoguchi, T., Shiba, T., Tamiaki, H., Kurisu, G., and Fujita, Y. (2010). X-ray crystal structure of the light-independent protochlorophyllide reductase. Nature 465, 110–114. 10.1038/nature08950.

34. Cheng, Q. (2008). Perspectives in biological nitrogen fixation research. J Integr Plant Biol 50, 786–798. 10.1111/j.1744-7909.2008.00700.x.

35. Fujita, Y., and Bauer, C.E. (2000). Reconstitution of light-independent protochlorophyllide reductase from purified bchl and BchN-BchB subunits. In vitro confirmation of nitrogenase-like features of a bacteriochlorophyll biosynthesis enzyme. J. Biol. Chem. 275, 23583–23588. 10.1074/jbc.M002904200.

36. Sarma, R., Barney, B.M., Hamilton, T.L., Jones, A., Seefeldt, L.C., and Peters, J.W. (2008). Crystal Structure of the L Protein of Rhodobacter Sphaeroides Light-Independent Protochlorophyllide Reductase with MgADP Bound: A Homologue of the Nitrogenase Fe Protein. Biochemistry 47, 13004–13015. 10.1021/bi801058r.

37. Nomata, J., Kitashima, M., Inoue, K., and Fujita, Y. (2006). Nitrogenase Fe protein-like Fe-S cluster is conserved in L-protein (BchL) of dark-operative protochlorophyllide reductase from Rhodobacter capsulatus. FEBS Lett. 580, 6151–6154. 10.1016/j.febslet.2006.10.014.

38. Cheng, Q., Day, A., Dowson-Day, M., Shen, G.F., and Dixon, R. (2005). The Klebsiella pneumoniae nitrogenase Fe protein gene (nifH) functionally substitutes for the chlL gene in Chlamydomonas reinhardtii. Biochem. Biophys. Res. Commun. 329, 966–975. 10.1016/j.bbrc.2005.02.064.

39. Reinbothe, C., El Bakkouri, M., Buhr, F., Muraki, N., Nomata, J., Kurisu, G., Fujita, Y., and Reinbothe, S. (2010). Chlorophyll biosynthesis: spotlight on protochlorophyllide reduction. Trends Plant Sci. 15, 614–624. 10.1016/j.tplants.2010.07.002.

40. Vedalankar, P., and Tripathy, B.C. (2023). Fortuitous events in the evolution of Light-dependent Protochlorophyllide Oxidoreductase. bioRxiv, 2023.2004. 2016.537069.

41. Chernomor, O., Peters, L., Schneidewind, J., Loeschcke, A., Knieps-Grunhagen, E., Schmitz, F., von Lieres, E., Kutta, R.J., Svensson, V., Jaeger, K.E., et al. (2021). Complex Evolution of Light-Dependent Protochlorophyllide Oxidoreductases in Aerobic Anoxygenic Phototrophs: Origin, Phylogeny, and Function. Mol. Biol. Evol. 38, 819–837. 10.1093/molbev/msaa234.

42. Soo, R.M., Hemp, J., Parks, D.H., Fischer, W.W., and Hugenholtz, P. (2017). On the origins of oxygenic photosynthesis and aerobic respiration in Cyanobacteria. Science 355, 1436–1440. 10.1126/science.aal3794.

43. Broecker, M.J., Schomburg, S., Heinz, D.W., Jahn, D., Schubert, W.D., and Moser, J. (2010). Crystal Structure of the Nitrogenase-like Dark Operative Protochlorophyllide Oxidoreductase Catalytic Complex (ChlN/ChlB)(2). J. Biol. Chem. 285, 27336–27345. 10.1074/jbc.M110.126698.

44. Dong, C.S., Zhang, W.L., Wang, Q., Li, Y.S., Wang, X., Zhang, M., and Liu, L. (2020). Crystal structures of cyanobacterial light-dependent protochlorophyllide oxidoreductase. P Natl Acad Sci USA 117, 8455–8461. 10.1073/pnas.1920244117.

45. Schneidewind, J., Krause, F., Bocola, M., Stadler, A.M., Davari, M.D., Schwaneberg, U., Jaeger, K.E., and Krauss, U. (2019). Consensus model of a cyanobacterial light-dependent protochlorophyllide oxidoreductase in its pigment-free apo-form and photoactive ternary complex. Commun Biol 2, 351. 10.1038/s42003-019-0590-4.

46. Boyd, E.S., and Peters, J.W. (2013). New insights into the evolutionary history of biological nitrogen fixation. Frontiers in microbiology 4, 201.

47. Layer, G., Krausze, J., and Moser, J. (2017). Reduction of Chemically Stable Multibonds: Nitrogenase-Like Biosynthesis of Tetrapyrroles. Adv. Exp. Med. Biol. 925, 147–161. 10.1007/5584_2016_175.

48. Hu, Y., and Ribbe, M.W. (2015). Nitrogenase and homologs. J Biol Inorg Chem 20, 435–445. 10.1007/s00775-014-1225-3.

49. Schmidt, F.V., Schulz, L., Zarzycki, J., Prinz, S., Oehlmann, N.N., Erb, T.J., and Rebelein, J.G. (2024). Structural insights into the iron nitrogenase complex. Nat. Struct. Mol. Biol. 31, 150–158.

50. Pereira-Leal, J.B., Levy, E.D., Kamp, C., and Teichmann, S.A. (2007). Evolution of protein complexes by duplication of homomeric interactions. Genome Biol 8, R51. 10.1186/gb-2007-8-4-r51.

51. Einsle, O., and Rees, D.C. (2020). Structural Enzymology of Nitrogenase Enzymes. Chem Rev 120, 4969–5004. 10.1021/acs.chemrev.0c00067.

52. van Kempen, M., Kim, S.S., Tumescheit, C., Mirdita, M., Lee, J., Gilchrist, C.L.M., Soding, J., and Steinegger, M. (2024). Fast and accurate protein structure search with Foldseek. Nat. Biotechnol. 42, 243–246. 10.1038/s41587-023-01773-0.

53. Endelman, J.B., Silberg, J.J., Wang, Z.G., and Arnold, F.H. (2004). Site-directed protein recombination as a shortest-path problem. Protein Engineering Design & Selection 17, 589–594. 10.1093/protein/gzh067.

54. Voigt, C.A., Martinez, C., Wang, Z.G., Mayo, S.L., and Arnold, F.H. (2002). Protein building blocks preserved by recombination. Nat. Struct. Biol. 9, 553–558. 10.1038/nsb805.

55. Buhr, F., El Bakkouri, M., Valdez, O., Pollmann, S., Lebedev, N., Reinbothe, S., and Reinbothe, C. (2008). Photoprotective role of NADPH: protochlorophyllide oxidoreductase A. P Natl Acad Sci USA 105, 12629–12634. 10.1073/pnas.0803950105.

56. Bi, C., Lu, N., Han, T., Huang, Z., Chen, J.Y., He, C., and Lu, Z. (2020). Whole-Genome Resequencing of Twenty Branchiostoma belcheri Individuals Provides a Brand-New Variant Dataset for Branchiostoma. Biomed Res Int 2020, 3697342. 10.1155/2020/3697342.

57. Guerardel, Y., Chang, L.Y., Fujita, A., Coddeville, B., Maes, E., Sato, C., Harduin-Lepers, A., Kubokawa, K., and Kitajima, K. (2012). Sialome analysis of the cephalochordate Branchiostoma belcheri, a key organism for vertebrate evolution. Glycobiology 22, 479–491. 10.1093/glycob/cwr155.

58. Longo, L.M., Jablonska, J., Vyas, P., Kanade, M., Kolodny, R., Ben-Tal, N., and Tawfik, D.S. (2020). On the emergence of P-Loop NTPase and Rossmann enzymes from a Beta-Alpha-Beta ancestral fragment. Elife 9. 10.7554/eLife.64415.

59. Imhoff, J.F., Rahn, T., Kunzel, S., and Neulinger, S.C. (2019). Phylogeny of Anoxygenic Photosynthesis Based on Sequences of Photosynthetic Reaction Center Proteins and a Key Enzyme in Bacteriochlorophyll Biosynthesis, the Chlorophyllide Reductase. Microorganisms 7. 10.3390/microorganisms7110576.

60. Nomata, J., Mizoguchi, T., Tamiaki, H., and Fujita, Y. (2006). A second nitrogenase-like enzyme for bacteriochlorophyll biosynthesis: reconstitution of chlorophyllide a reductase with purified X-protein (BchX) and YZ-protein (BchY-BchZ) from Rhodobacter capsulatus. J. Biol. Chem. 281, 15021–15028. 10.1074/jbc.M601750200.

61. Sahu, B., and Maeda, A. (2016). Retinol Dehydrogenases Regulate Vitamin A Metabolism for Visual Function. Nutrients 8. 10.3390/nu8110746.

62. Lhor, M., and Salesse, C. (2014). Retinol dehydrogenases: membrane-bound enzymes for the visual function. Biochem. Cell Biol. 92, 510–523. 10.1139/bcb-2014-0082.

63. Imasheva, E.S., Balashov, S.P., Choi, A.R., Jung, K.H., and Lanyi, J.K. (2009). Reconstitution of Gloeobacter violaceus rhodopsin with a light-harvesting carotenoid antenna. Biochemistry 48, 10948–10955. 10.1021/bi901552x.

64. Chazan, A., Das, I., Fujiwara, T., Murakoshi, S., Rozenberg, A., Molina-Marquez, A., Sano, F.K., Tanaka, T., Gomez-Villegas, P., Larom, S., et al. (2023). Phototrophy by antenna-containing rhodopsin pumps in aquatic environments. Nature 615, 535–540. 10.1038/s41586-023-05774-6.

65. Oesterhelt, D., and Stoeckenius, W. (1973). Functions of a new photoreceptor membrane. Proc Natl Acad Sci U S A 70, 2853–2857. 10.1073/pnas.70.10.2853.

66. Balashov, S.P., Imasheva, E.S., Boichenko, V.A., Anton, J., Wang, J.M., and Lanyi, J.K. (2005). Xanthorhodopsin: a proton pump with a light-harvesting carotenoid antenna. Science 309, 2061–2064. 10.1126/science.1118046.

67. Kojima, K., and Sudo, Y. (2023). Convergent evolution of animal and microbial rhodopsins. RSC Adv 13, 5367–5381. 10.1039/d2ra07073a.

68. Chen, J.H., Wu, H., Xu, C., Liu, X.C., Huang, Z., Chang, S., Wang, W., Han, G., Kuang, T., Shen, J.R., and Zhang, X. (2020). Architecture of the photosynthetic complex from a green sulfur bacterium. Science 370. 10.1126/science.abb6350.

69. Tsuji, J.M., Shaw, N.A., Nagashima, S., Venkiteswaran, J.J., Schiff, S.L., Watanabe, T., Fukui, M., Hanada, S., Tank, M., and Neufeld, J.D. (2024). Anoxygenic phototroph of the Chloroflexota uses a type I reaction centre. Nature 627, 915–922. 10.1038/s41586-024-07180-y.

70. Banda, D.M., Pereira, J.H., Liu, A.K., Orr, D.J., Hammel, M., He, C., Parry, M.A.J., Carmo-Silva, E., Adams, P.D., Banfield, J.F., and Shih, P.M. (2020). Novel bacterial clade reveals origin of form I Rubisco. Nat Plants 6, 1158–1166. 10.1038/s41477-020-00762-4.

71. Oliver, N., Avramov, A.P., Nurnberg, D.J., Dau, H., and Burnap, R.L. (2022). From manganese oxidation to water oxidation: assembly and evolution of the water-splitting complex in photosystem II. Photosynth Res 152, 107–133. 10.1007/s11120-022-00912-z.

72. Cardona, T. (2018). Early Archean origin of heterodimeric Photosystem I. Heliyon 4, e00548. 10.1016/j.heliyon.2018.e00548.

73. Rutherford, A.W., and Faller, P. (2003). Photosystem II: evolutionary perspectives. Philos Trans R Soc Lond B Biol Sci 358, 245–253. 10.1098/rstb.2002.1186.

74. Nelson, N., and Ben-Shem, A. (2005). The structure of photosystem I and evolution of photosynthesis. Bioessays 27, 914–922. 10.1002/bies.20278.

75. Cardona, T., Murray, J.W., and Rutherford, A.W. (2015). Origin and Evolution of Water Oxidation before the Last Common Ancestor of the Cyanobacteria. Mol. Biol. Evol. 32, 1310–1328. 10.1093/molbev/msv024.

76. Silva, P.J., and Cheng, Q. (2022). An Alternative Proposal for the Reaction Mechanism of Light-Dependent Protochlorophyllide Oxidoreductase. Acs Catalysis 12, 2589–2605. 10.1021/acscatal.1c05351.

77. Cheng, Q. (2013). The Earth is a Clock: A Natural History of Biological Nitrogen Fixation and the Planetary Future (Science Press ISBN978-7-03-038773-8).

78. Xu, H.F., Yu, C., Bai, Y., Zuo, A.W., Ye, Y.T., Liu, Y.R., Li, Z.K., Dai, G.Z., Chen, M., and Qiu, B.S. (2024). Red-light-dependent chlorophyll synthesis kindles photosynthetic recovery of chlorotic dormant cyanobacteria using a dark-operative enzyme. Curr. Biol. 34, 4424–4435 e4423. 10.1016/j.cub.2024.07.083.

79. Allen, J.F., Thake, B., and Martin, W.F. (2019). Nitrogenase Inhibition Limited Oxygenation of Earth’s Proterozoic Atmosphere. Trends Plant Sci. 24, 1022–1031. 10.1016/j.tplants.2019.07.007.

80. Xun, H., Wang, Y., Yuan, J., Lian, L., Feng, W., Liu, S., Hong, J., Liu, B., Ma, J., and Wang, X. (2024). Non-CG DNA hypomethylation promotes photosynthesis and nitrogen fixation in soybean. Proc Natl Acad Sci U S A 121, e2402946121. 10.1073/pnas.2402946121.

81. Land, M.F., and Nilsson, D.-E. (2012). viiPreface to the second edition. In Animal Eyes, (Oxford University Press), pp. 0. 10.1093/acprof:oso/9780199581139.002.0006.

82. Gehring, W.J. (2014). The evolution of vision. Wiley Interdiscip Rev Dev Biol 3, 1–40. 10.1002/wdev.96.

83. Lamb, T.D., Collin, S.P., and Pugh, E.N., Jr. (2007). Evolution of the vertebrate eye: opsins, photoreceptors, retina and eye cup. Nat. Rev. Neurosci. 8, 960–976. 10.1038/nrn2283.

84. Nilsson, D.E. (2013). Eye evolution and its functional basis. Vis Neurosci 30, 5–20. 10.1017/S0952523813000035.

85. Blankenship, R.E. (2010). Early evolution of photosynthesis. Plant Physiol. 154, 434–438. 10.1104/pp.110.161687.

86. Knoll, A.H. (2015). Life on a Young Planet: The First Three Billion Years of Evolution on Earth (Princeton University Press).

87. Heyes, D.J., Heathcote, P., Rigby, S.E., Palacios, M.A., van Grondelle, R., and Hunter, C.N. (2006). The first catalytic step of the light-driven enzyme protochlorophyllide oxidoreductase proceeds via a charge transfer complex. J. Biol. Chem. 281, 26847–26853. 10.1074/jbc.M602943200.

88. Heyes, D.J., Hunter, C.N., van Stokkum, I.H., Van Grondelle, R., and Groot, M.L. (2003). Ultrafast enzymatic reaction dynamics in protochlorophyllide oxidoreductase. Nat. Struct. Mol. Biol. 10, 491–492.

89. Weiss, M.C., Preiner, M., Xavier, J.C., Zimorski, V., and Martin, W.F. (2018). The last universal common ancestor between ancient Earth chemistry and the onset of genetics. PLoS Genet. 14, e1007518.

90. Martin, W., and Russell, M.J. (2003). On the origins of cells: a hypothesis for the evolutionary transitions from abiotic geochemistry to chemoautotrophic prokaryotes, and from prokaryotes to nucleated cells. Philos Trans R Soc Lond B Biol Sci 358, 59–83; discussion 83-55. 10.1098/rstb.2002.1183.

91. Ohki, Y., Munakata, K., Matsuoka, Y., Hara, R., Kachi, M., Uchida, K., Tada, M., Cramer, R.E., Sameera, W.M.C., Takayama, T., et al. (2022). Nitrogen reduction by the Fe sites of synthetic [Mo3S4Fe] cubes. Nature 607, 86-+. 10.1038/s41586-022-04848-1.

92. Jeoung, J.H., Martins, B.M., and Dobbek, H. (2020). Double-Cubane [8Fe9S] Clusters: A Novel Nitrogenase-Related Cofactor in Biology. ChemBioChem 21, 1710–1716. 10.1002/cbic.202000016.

93. Reardon, S. (2024). How synthetic biologists are building better biofactories. Nature 628, 224–226. 10.1038/d41586-024-00907-x.

94. Edman, N.I., Phal, A., Redler, R.L., Schlichthaerle, T., Srivatsan, S.R., Ehnes, D.D., Etemadi, A., An, S.J., Favor, A., Li, Z., et al. (2024). Modulation of FGF pathway signaling and vascular differentiation using designed oligomeric assemblies. Cell 187, 3726–3740 e3743. 10.1016/j.cell.2024.05.025.

95. Huang, B., Abedi, M., Ahn, G., Coventry, B., Sappington, I., Tang, C., Wang, R., Schlichthaerle, T., Zhang, J.Z., Wang, Y., et al. (2024). Designed endocytosis-inducing proteins degrade targets and amplify signals. Nature. 10.1038/s41586-024-07948-2.

96. Jumper, J., Evans, R., Pritzel, A., Green, T., Figurnov, M., Ronneberger, O., Tunyasuvunakool, K., Bates, R., Zidek, A., Potapenko, A., et al. (2021). Highly accurate protein structure prediction with AlphaFold. Nature 596, 583–589. 10.1038/s41586-021-03819-2.

97. Almagro Armenteros, J.J., Salvatore, M., Emanuelsson, O., Winther, O., von Heijne, G., Elofsson, A., and Nielsen, H. (2019). Detecting sequence signals in targeting peptides using deep learning. Life Sci Alliance 2. 10.26508/lsa.201900429.

98. Stivala, A., Wybrow, M., Wirth, A., Whisstock, J.C., and Stuckey, P.J. (2011). Automatic generation of protein structure cartoons with Pro-origami. Bioinformatics 27, 3315–3316. 10.1093/bioinformatics/btr575.

99. Holm, L. (2022). Dali server: structural unification of protein families. Nucleic Acids Res. 50, W210–W215. 10.1093/nar/gkac387.

100. Chen, C., Chen, H., Zhang, Y., Thomas, H.R., Frank, M.H., He, Y., and Xia, R. (2020). TBtools: An Integrative Toolkit Developed for Interactive Analyses of Big Biological Data. Mol Plant 13, 1194–1202. 10.1016/j.molp.2020.06.009.

101. Zhang, C., Shine, M., Pyle, A.M., and Zhang, Y. (2022). US-align: universal structure alignments of proteins, nucleic acids, and macromolecular complexes. Nat. Methods 19, 1109–1115. 10.1038/s41592-022-01585-1.

102. Trott, O., and Olson, A.J. (2010). AutoDock Vina: improving the speed and accuracy of docking with a new scoring function, efficient optimization, and multithreading. J Comput Chem 31, 455–461. 10.1002/jcc.21334.

103. Abramson, J., Adler, J., Dunger, J., Evans, R., Green, T., Pritzel, A., Ronneberger, O., Willmore, L., Ballard, A.J., Bambrick, J., et al. (2024). Accurate structure prediction of biomolecular interactions with AlphaFold 3. Nature 630, 493–500. 10.1038/s41586-024-07487-w.

104. Krieger, E., and Vriend, G. (2015). New ways to boost molecular dynamics simulations. J Comput Chem 36, 996–1007. 10.1002/jcc.23899.

105. Jakalian, A., Jack, D.B., and Bayly, C.I. (2002). Fast, efficient generation of high-quality atomic charges. AM1-BCC model: II. Parameterization and validation. J Comput Chem 23, 1623–1641. 10.1002/jcc.10128.

106. Neese, F. (2022). Software update: The ORCA program system-Version 5.0. Wires Comput Mol Sci 12. ARTN e160610.1002/wcms.1606.

107. Bayly, C.I., Cieplak, P., Cornell, W., and Kollman, P.A. (1993). A well-behaved electrostatic potential based method using charge restraints for deriving atomic charges: the RESP model. The Journal of Physical Chemistry 97, 10269–10280.

108. Lu, T., and Chen, F. (2012). Multiwfn: a multifunctional wavefunction analyzer. J Comput Chem 33, 580–592. 10.1002/jcc.22885.

109. Essmann, U., Perera, L., Berkowitz, M.L., Darden, T., Lee, H., and Pedersen, L.G. (1995). A Smooth Particle Mesh Ewald Method. J Chem Phys 103, 8577–8593. Doi 10.1063/1.470117.

110. Kumar, S., Rosenberg, J.M., Bouzida, D., Swendsen, R.H., and Kollman, P.A. (1992). The weighted histogram analysis method for free-energy calculations on biomolecules. I. The method. Journal of computational chemistry 13, 1011–1021.

