## Supplemental figure for "Structure-based analysis unveils co-origin of LPOR and nitrogenase-like proteins"

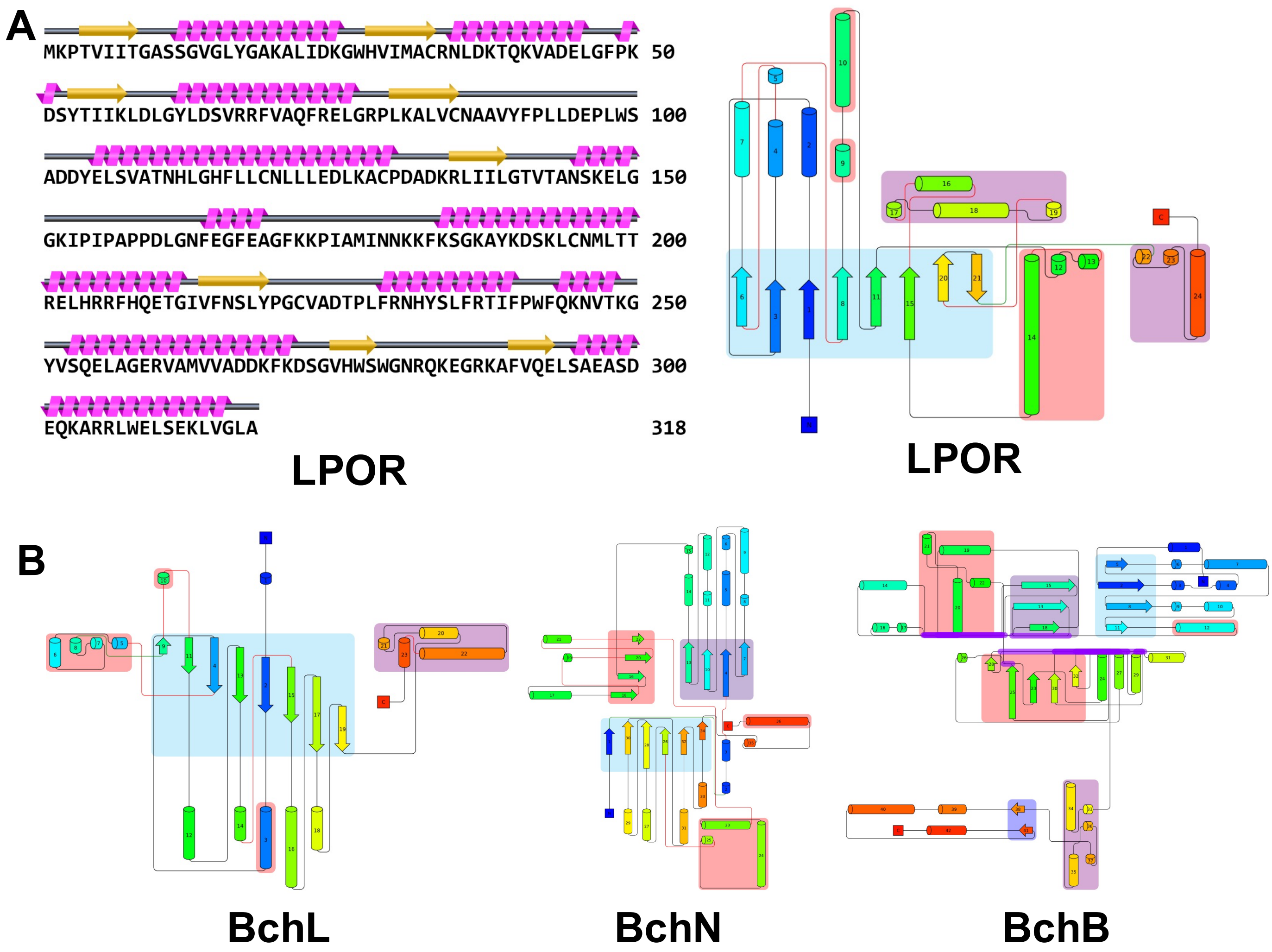


**Figure S1. The secondary structure topology map of LPOR and different subunits of DPOR.** Topological graphs of LPOR **(A)** and different subunits of DPOR **(B)** were generated using Pro-origami. The graph secondary structure elements are represented and color-coded from the N-terminal (blue) to the C-terminal (red). The shaded areas indicate the core domains as predicted by Pro-origami.

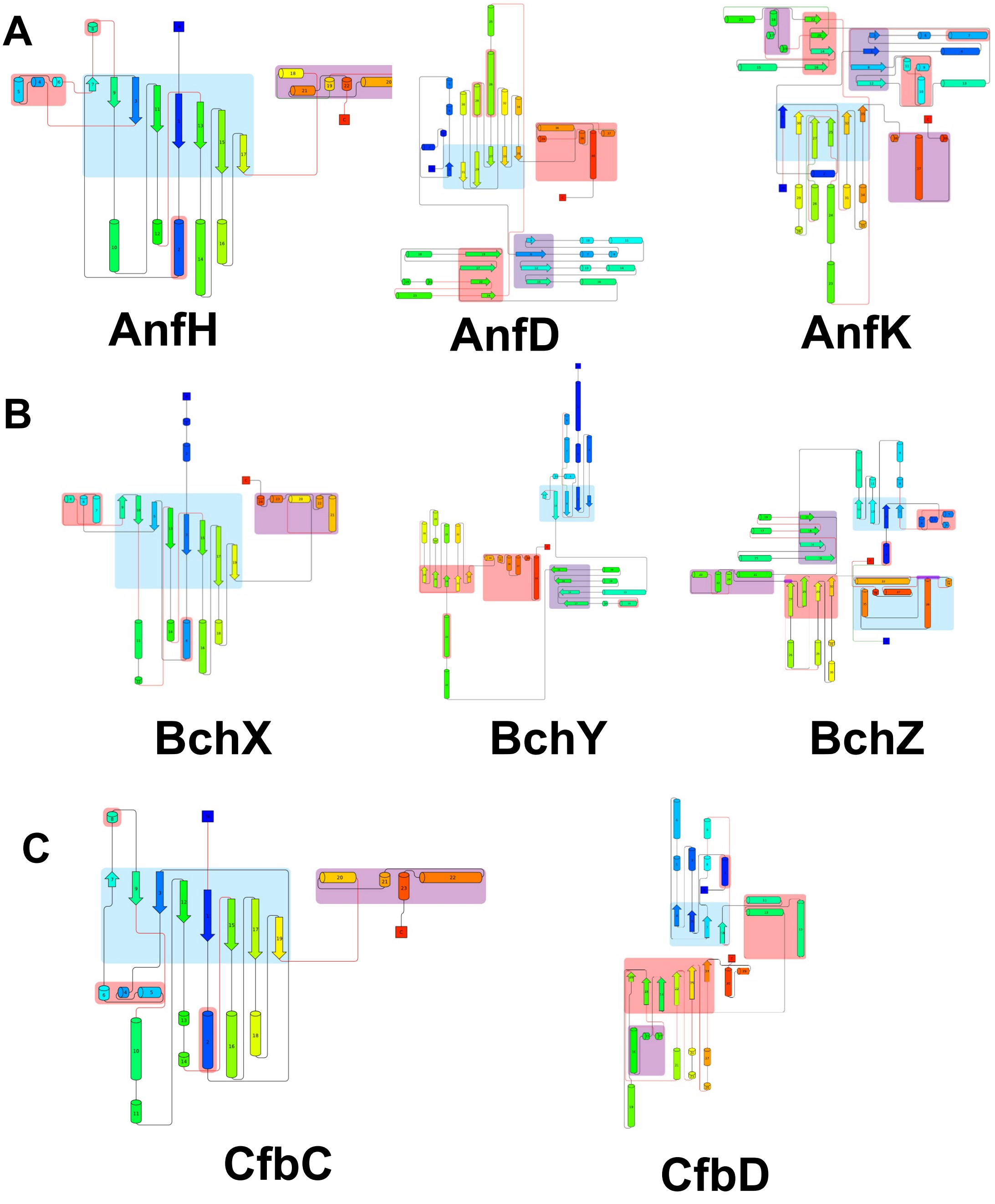


**Figure S2. The secondary structure topology map of different subunits of nitrogenase, COR and F430.** Topological graphs of LPOR **(A)** and different subunits of nitrogenase **(A)**, COR **(B)** and F430 **(C)** were generated using Pro-origami. The graph secondary structure elements are represented and color-coded from the N-terminal (blue) to the C-terminal (red). The shaded areas indicate the core domains as predicted by Pro-origami.

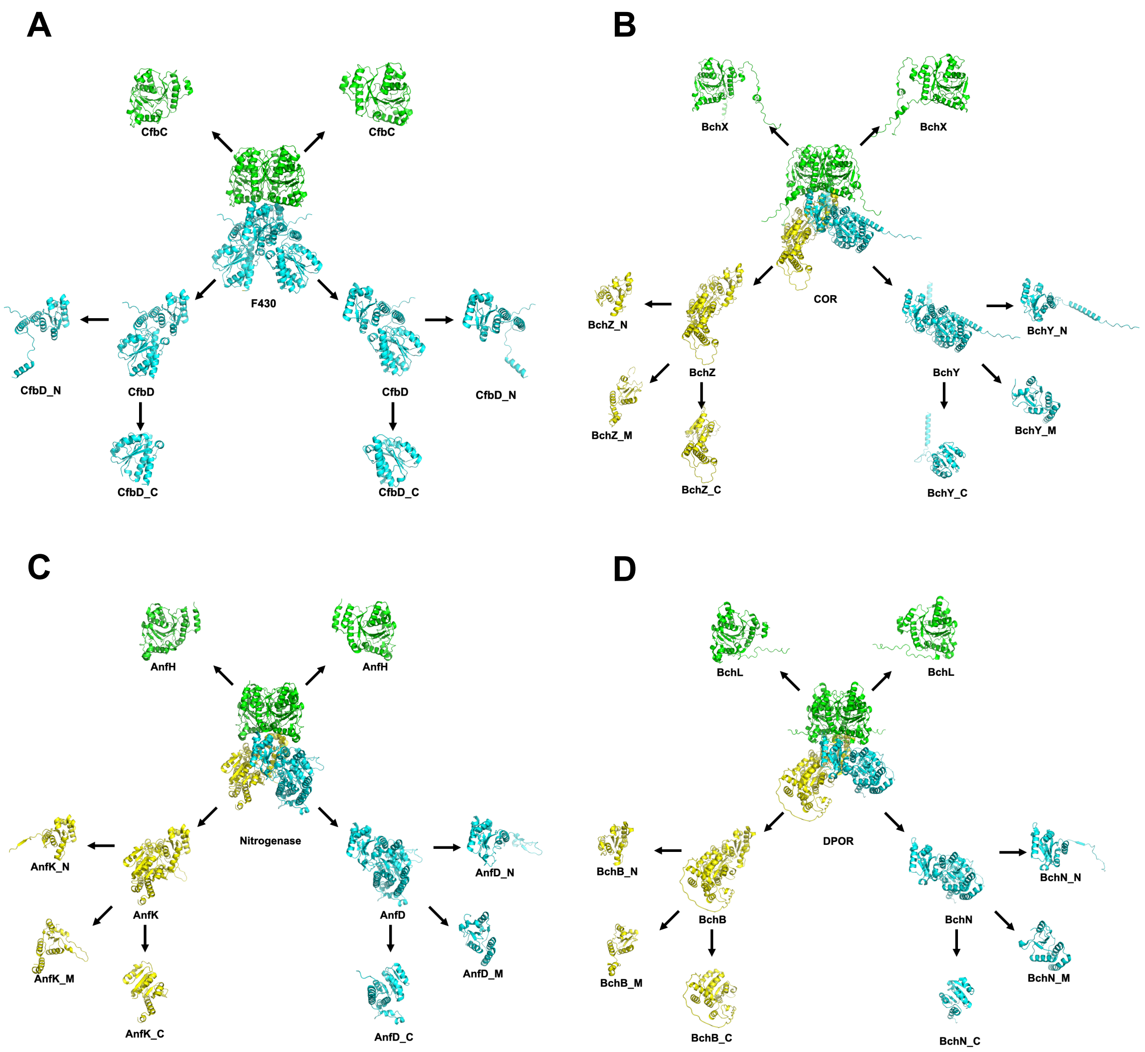


**Figure S3. The ribbon diagrams of full structures, different subunits and split individual subdomains of F430 (A), COR (B), nitrogenase (C) and DPOR (D).**

**AFDB-SWISSPROT** 39 hits

GRAPHICAL | NUMERIC

| Target | Description | Scientific Name | Prob. | Seq. Id. | TM-score | Position in query | Alignment |
|---|---|---|---|---|---|---|---|
| AF-Q8TJZ8-F1-model_v4 | Ni-sirohydrochlorin a,c-diami… | Methanosarcina acetivorans … | 1.00 | 100 | 0.756 | 1 – 196 | |
| AF-Q8TPT7-F1-model_v4 | 6,7-dimethyl-8-ribityllumazin… | Methanosarcina acetivorans … | 0.18 | 6.2 | 0.447 | 26 – 146 | |
| AF-Q8TIP4-F1-model_v4 | UPF0292 protein MA_4098 | Methanosarcina acetivorans … | 0.18 | 3.8 | 0.434 | 3 – 180 | |
| AF-Q8TSS3-F1-model_v4 | O-phospho-L-seryl-tRNA:Cy… | Methanosarcina acetivorans … | 0.57 | 6.6 | 0.422 | 1 – 167 | |
| AF-Q8TJZ9-F1-model_v4 | Ni-sirohydrochlorin a,c-diami… | Methanosarcina acetivorans … | 0.30 | 5.1 | 0.398 | 12 – 170 | |
| AF-Q8TK94-F1-model_v4 | Serine hydroxymethyltransfe… | Methanosarcina acetivorans … | 0.44 | 5.1 | 0.385 | 2 – 166 | |
| AF-P58831-F1-model_v4 | UPF0200 protein MA_4660 | Methanosarcina acetivorans … | 0.16 | 5 | 0.383 | 13 – 146 | |
| AF-Q8THC3-F1-model_v4 | Shikimate dehydrogenase (… | Methanosarcina acetivorans … | 0.28 | 4.2 | 0.379 | 1 – 172 | |
| AF-Q8TUE9-F1-model_v4 | Histidinol-phosphate aminotr… | Methanosarcina acetivorans … | 0.38 | 3.2 | 0.378 | 7 – 166 | |
| AF-Q8TRA5-F1-model_v4 | S-inosyl-L-homocysteine hyd… | Methanosarcina acetivorans … | 0.41 | 4.5 | 0.372 | 1 – 181 | |

**Figure S4. Structural homology search of CfbD_N using Foldseek.** CfbD_N was used as an input to search the Swiss-Prot database for *Methanosarcina acetivorans*. The fourth-ranked match (highlighted in red box) corresponds to Ni-sirohydrochlorin a,c-diamide synthase (CfbC), with a TM-score of 0.398 and 5.1% sequence identity.

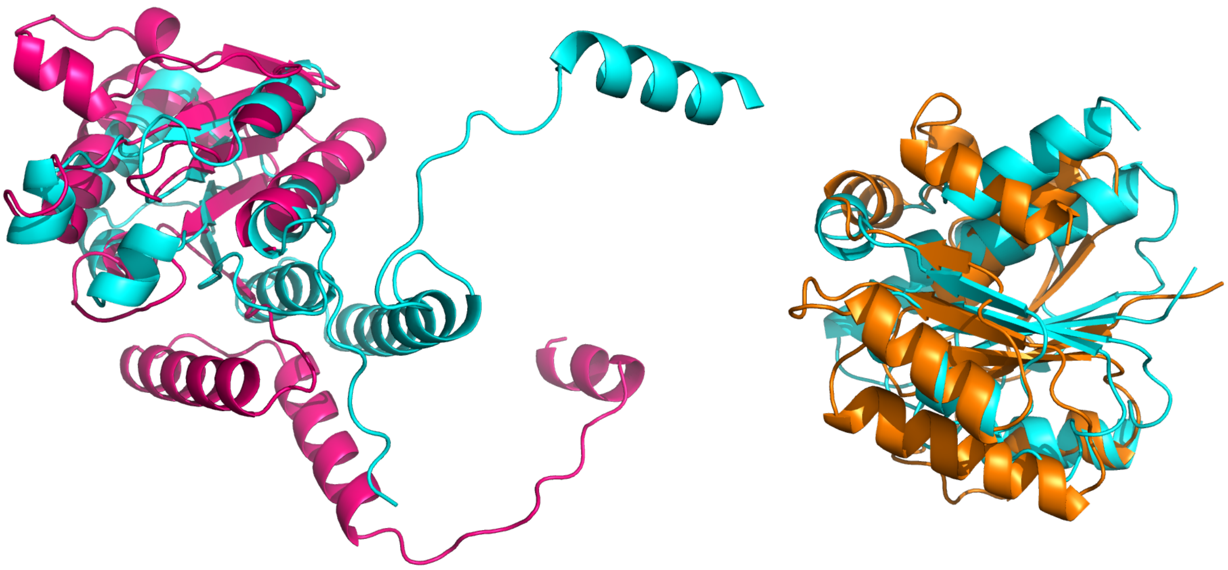

**CfbD_N vs Simulated CfbD_N**
**Z-score=5.6**

**CfbD_C vs Simulated CfbD_C**
**Z-score=4.7**

**Figure S5. The structural alignments of AlphaFold2-prdicted structures of assembled CfbD_N and CfbD_C with AlphaFold2-predicted target structures of CfbD_N and CfbD_C.** The assembled CfbD_N and CfbD_C were generated by extracting the corresponding amino acid sequences of CfbC showing structural similarity with CfbD. Structural alignments of the target CfbD with the assembled CfbD_N and CfbD_C in ribbon representation using US-align, respectively. The target structure of CfbD is in green, and the structures of assembled CfbD_N and CfbD_C are in pink and orange, respectively. The structural similarity threshold is DALI Z-score of 2. All the structures were predicted by AlphaFold2.

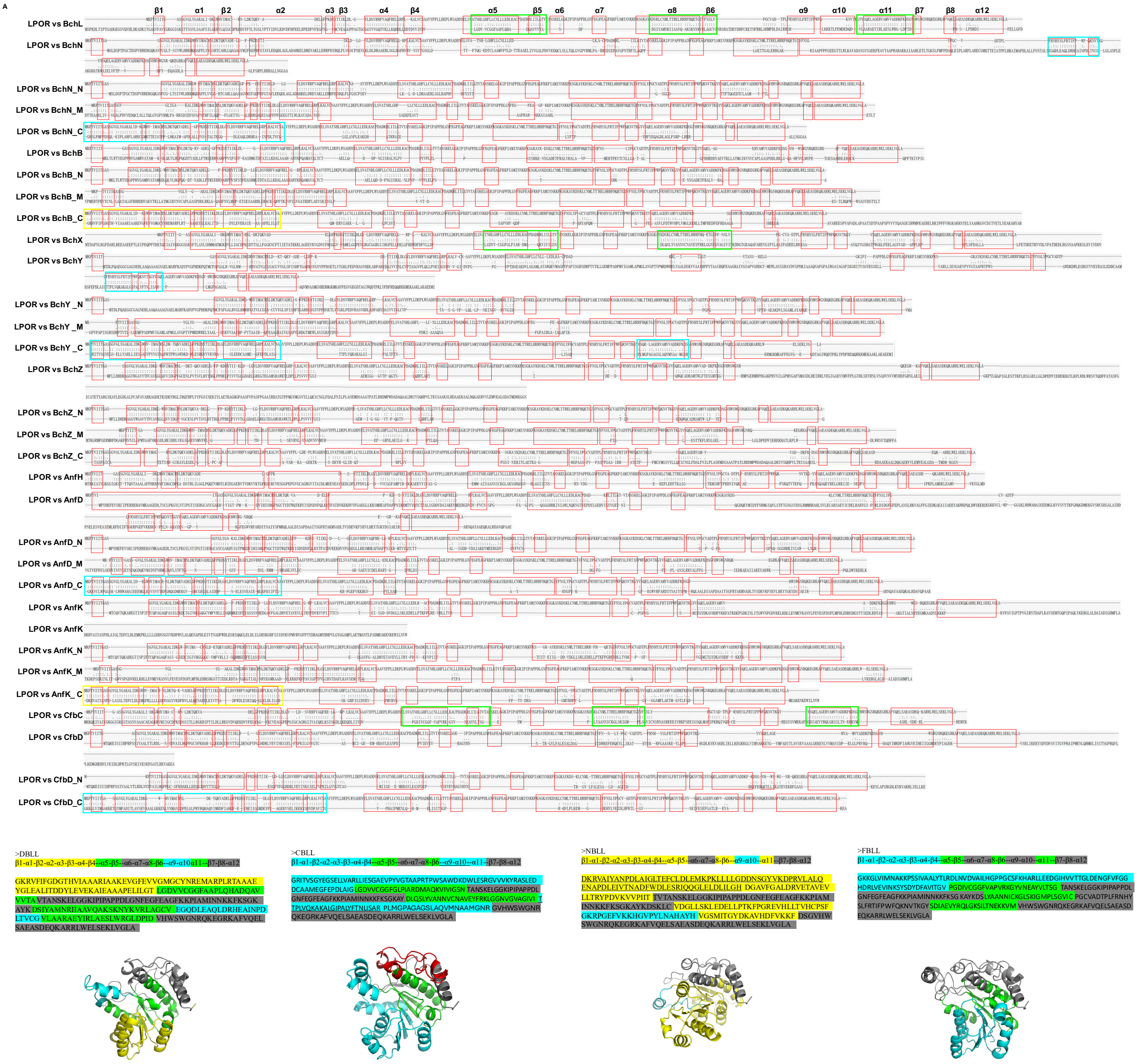


**Figure S6. Design of non-natural LPOR analogs (nnLPORs) by assembling similar amino acid sequences based on the structural similarities**. **A**). Secondary structure alignments between LPOR and selected subunits from DPOR, COR, nitrogenase, and the F430 complex, were conducted using the US-align tool. The alignments highlight amino acid fragments showing significant structural similarities, which were subsequently selected to assemble the nnLPOR amino acid sequences. ":" denotes residue pairs of d < 5.0 Angstrom, "." denotes other aligned residues. **B**). Schematic representations of the assembled nnLPOR amino acid sequences. Structurally conserved amino acid fragments identified from the comparative structure analysis were integrated and highlighted with different colors. **C**). Ribbon diagrams of the AlphaFold2-predicted nnLPORs structures. Three-dimensional structural predictions of nnLPORs, including DBLL, CBLL, NBLL and FBLL, by AlphaFold2, demonstrating substantial structural alignment with the archetype LPOR enzyme.

**AFDB-SWISSPROT** 867 hits

GRAPHICAL NUMERIC

| Target | Description | Scientific Name | Prob. | Seq. Id. | E-Value | Position in query | Alignment |
|---|---|---|---|---|---|---|---|
| AF-Q59987-F1-model_v4 | Light-dependent protochloro… | Synechocystis sp. PCC 680… | 1.00 | 96.5 | 2.39e-56 | 1–318 | |
| AF-O66148-F1-model_v4 | Light-dependent protochloro… | Leptolyngbya boryana | 1.00 | 74.2 | 1.31e-49 | 1–318 | |
| AF-Q39617-F1-model_v4 | Protochlorophyllide reductas… | Chlamydomonas reinhardtii | 1.00 | 52 | 5.00e-40 | 1–318 | |
| AF-O80333-F1-model_v4 | Protochlorophyllide reductas… | Marchantia paleacea | 1.00 | 55.7 | 1.29e-40 | 1–318 | |
| AF-O48741-F1-model_v4 | Protochlorophyllide reductas… | Arabidopsis thaliana | 1.00 | 54.8 | 1.08e-39 | 2–318 | |
| AF-Q41249-F1-model_v4 | Protochlorophyllide reductas… | Cucumis sativus | 1.00 | 54.8 | 4.18e-39 | 2–318 | |
| AF-P21218-F1-model_v4 | Protochlorophyllide reductas… | Arabidopsis thaliana | 1.00 | 55.3 | 3.11e-39 | 2–318 | |
| AF-Q9SDT1-F1-model_v4 | Protochlorophyllide reductas… | Daucus carota | 1.00 | 55.6 | 6.32e-39 | 2–318 | |
| AF-Q01289-F1-model_v4 | Protochlorophyllide reductas… | Pisum sativum | 1.00 | 55 | 1.21e-38 | 2–318 | |
| AF-Q42850-F1-model_v4 | Protochlorophyllide reductas… | Hordeum vulgare | 1.00 | 55.3 | 4.70e-38 | 2–318 | |
| AF-Q42536-F1-model_v4 | Protochlorophyllide reductas… | Arabidopsis thaliana | 1.00 | 54.6 | 1.01e-37 | 2–318 | |
| AF-Q7XKF3-F1-model_v4 | Protochlorophyllide reductas… | Oryza sativa Japonica Group | 1.00 | 53.2 | 2.60e-37 | 2–318 | |
| AF-Q8W3D9-F1-model_v4 | Protochlorophyllide reductas… | Oryza sativa Japonica Group | 1.00 | 52.8 | 1.44e-37 | 2–318 | |
| AF-P15904-F1-model_v4 | Protochlorophyllide reductase | Avena sativa | 1.00 | 52.2 | 1.36e-37 | 5–318 | |
| AF-P13653-F1-model_v4 | Protochlorophyllide reductas… | Hordeum vulgare | 1.00 | 52.9 | 2.92e-37 | 2–318 | |
| AF-Q41578-F1-model_v4 | Protochlorophyllide reductas… | Triticum aestivum | 1.00 | 52.3 | 5.27e-37 | 2–318 | |
| AF-Q8NBN7-F1-model_v4 | Retinol dehydrogenase 13 | Homo sapiens | 1.00 | 31.2 | 6.90e-22 | 3–318 | |
| AF-A2RVM0-F1-model_v4 | Short-chain dehydrogenase … | Arabidopsis thaliana | 1.00 | 27.3 | 4.30e-21 | 3–316 | |
| AF-Q8CEE7-F1-model_v4 | Retinol dehydrogenase 13 | Mus musculus | 1.00 | 32.7 | 3.20e-21 | 3–318 | |
| AF-Q6RVV4-F1-model_v4 | Short-chain dehydrogenase … | Pisum sativum | 1.00 | 29 | 1.04e-20 | 3–318 | |

**Figure S7. Structural homology search of LPOR using Foldseek.** The structure of LPOR was used as a query to search the AlphaFold2 Swiss-Prot database using Foldseek. Excluding homologous LPOR proteins, the top hit (highlighted in red box) is human retinol dehydrogenase 13 (RDH13), with a sequence identity of 31.2% and a TM-score of 0.792.

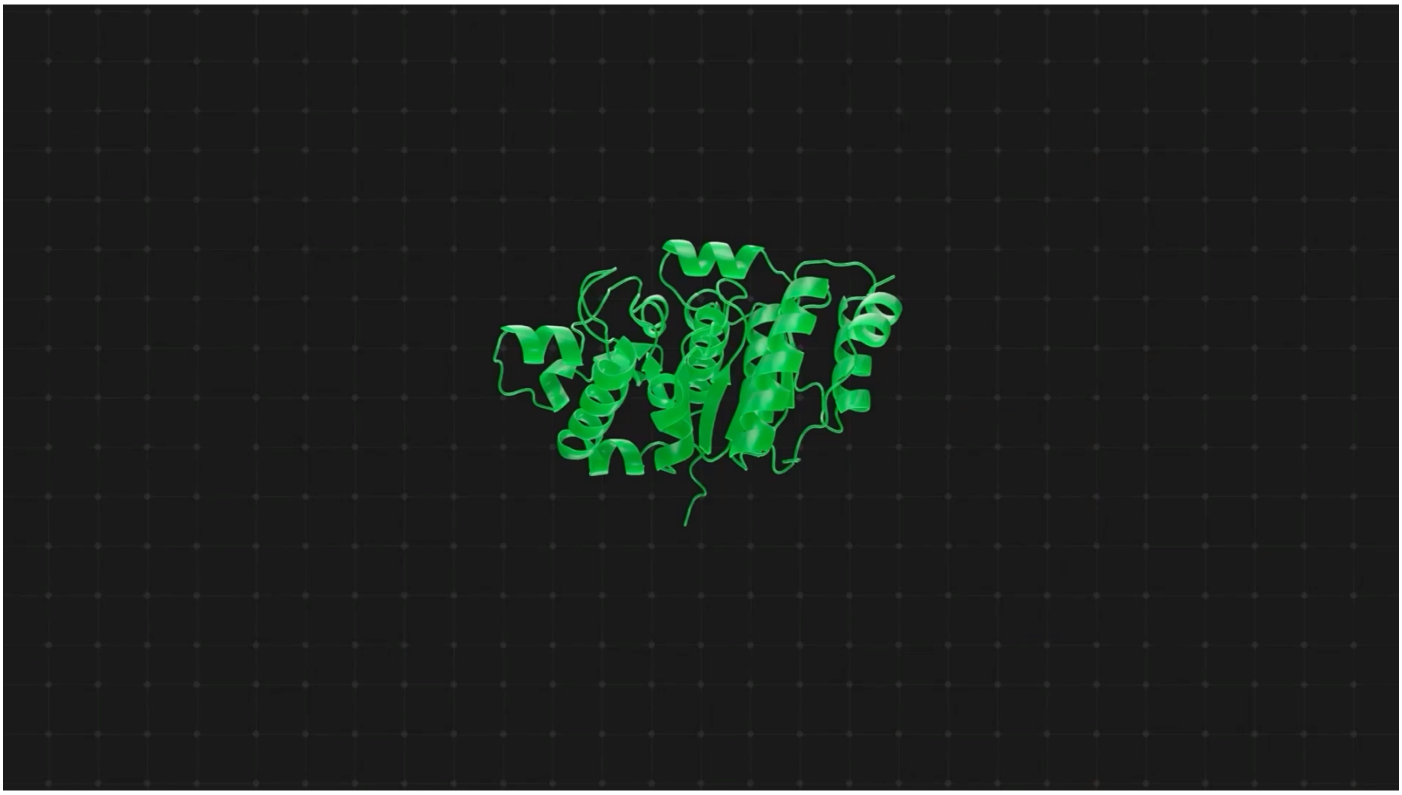

**Video S1. Structural origins and evolution of nitrogenase-like proteins, related to Figure 5**

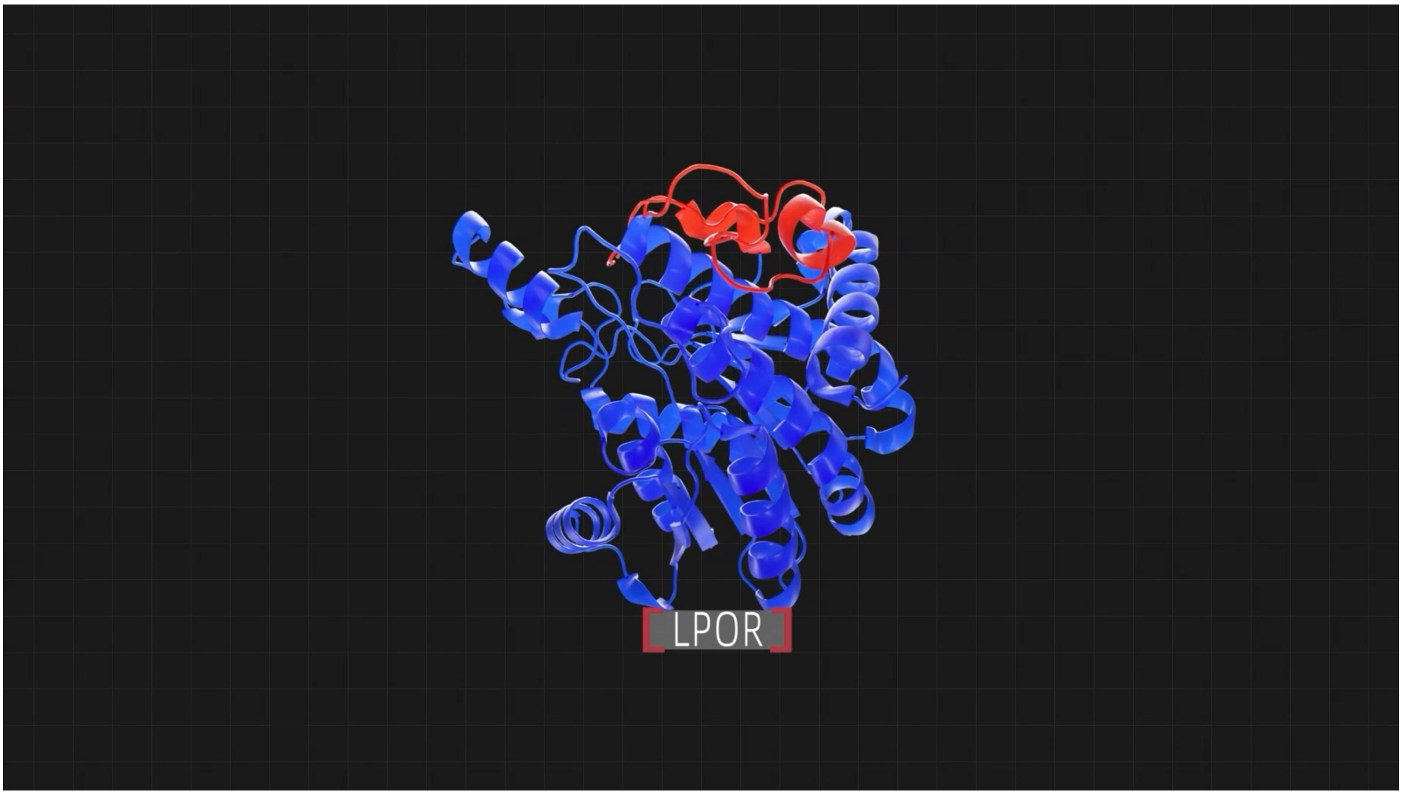


**Video S2. Structural origins and evolution of LPOR, related to Figure 5**

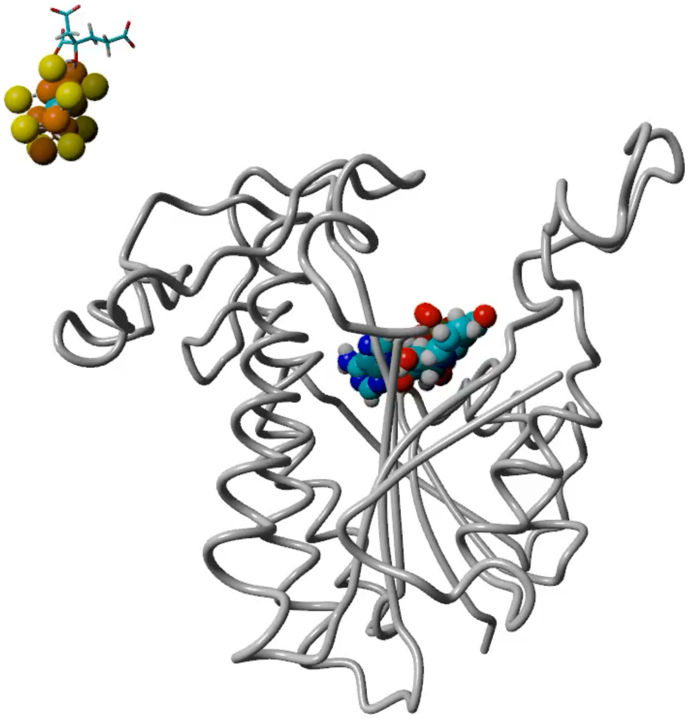

**Video S3. FeMoco integration into the engineered LPOR, related to Figure 7F**

The engineered LPOR protein structure was rendered as a pipe model using YASARA to clearly visualize ligand movement. The Fe and S atoms in FeMoco are depicted as orange and yellow spheres, respectively. HCA is depicted as a stick model positioned close to FeMoco. NADPH is depicted as solid space-filling sphere.
